# Revealing metabolic flexibility of *Candidatus* Accumulibacter phosphatis through redox cofactor analysis and metabolic network modeling

**DOI:** 10.1101/458331

**Authors:** Leonor Guedes da Silva, Karel Olavarria Gamez, Joana Castro Gomes, Kasper Akkermans, Laurens Welles, Ben Abbas, Mark C.M. van Loosdrecht, Sebastian Aljoscha Wahl

**Author notes:** Address correspondence to Leonor Guedes da Silva.

## Abstract

Environmental fluctuations in the availability of nutrients lead to intricate metabolic strategies. *Candidatus* Accumulibacter phosphatis, a polyphosphate accumulating organism (PAO) responsible for enhanced biological phosphorus removal (EBPR) from wastewater treatment systems, is prevalent in aerobic/anaerobic environments. While the overall metabolic traits of these bacteria are well described, the inexistence of isolates has led to controversial conclusions on the metabolic pathways used.

Here, we experimentally determined the redox cofactor preference of different oxidoreductases in the central carbon metabolism of a highly enriched *Ca*. A. phosphatis culture. Remarkably, we observed that the acetoacetyl-CoA reductase engaged in polyhydroxyalkanoates (PHA) synthesis is NADH-preferring instead of the generally assumed NADPH dependency. Based on previously published meta-omics data and the results of enzymatic assays, a reduced central carbon metabolic network was constructed and used for simulating different metabolic operating modes. In particular, scenarios with different acetate-to-glycogen consumption ratios were simulated. For a high ratio (i.e. more acetate), a polyphosphate-based metabolism arises as optimal with a metabolic flux through the glyoxylate shunt. In case of a low acetate-to-glycogen ratio, glycolysis is used in combination with reductive branch of the TCA cycle. Thus, optimal metabolic flux strategies will depend on the environment (acetate uptake) and on intracellular storage compounds availability (polyphosphate/glycogen).

This metabolic flexibility is enabled by the NADH-driven PHA synthesis. It allows for maintaining metabolic activity under varying environmental substrate conditions, with high carbon conservation and lower energetic costs compared to NADPH dependent PHA synthesis. Such (flexible) metabolic redox coupling can explain PAOs’ competitiveness under oxygen-fluctuating environments.

**IMPORTANCE:** Here we demonstrate how microbial metabolism can adjust to a wide range of environmental conditions. Such flexibility generates a selective advantage under fluctuating environmental conditions. It can also explain the different observations reported in PAO literature, including the capacity of *Ca*. Accumulibacter phosphatis to act like glycogen accumulating organisms (GAO). These observations stem from slightly different experimental conditions and controversy only arises when one assumes metabolism can only operate in one single mode. Furthermore, we also show how the study of metabolic strategies is possible when combining-omics data with functional assays and modeling. Genomic information can only provide the potential of a microorganism. The environmental context and other complementary approaches are still needed to study and predict the functional application of such metabolic potential.

## INTRODUCTION

Natural habitats of microorganisms are dynamic environments with intermittent supply of nutrients. Under these dynamic conditions, organisms are selected that can accumulate growth substrates when these are abundant to compensate for periods when these are exhausted (1).

Enhanced biological phosphate removal (EBPR) from wastewater is designed to make use of such physiological feature by circulating activated sludge through alternating zones with or without an external electron acceptor here respectively defined as aerobic/anaerobic (see Figure 1A) (4, 5). This environment selects for polyphosphate accumulating organisms (PAOs) like *Candidatus* Accumulibacter phosphatis (hereafter referred to as Accumulibacter). These bacteria thrive under these dynamic conditions thanks to a complex metabolic strategy encompassing the cycling of three common storage polymers: polyphosphate, glycogen and polyhydroxyalkanoates (PHA). Of these, polyphosphate stands out for allowing for fast and competitive harvesting of organic matter in the absence of an external electron acceptor (see Figure 1B).

**Figure 1.**
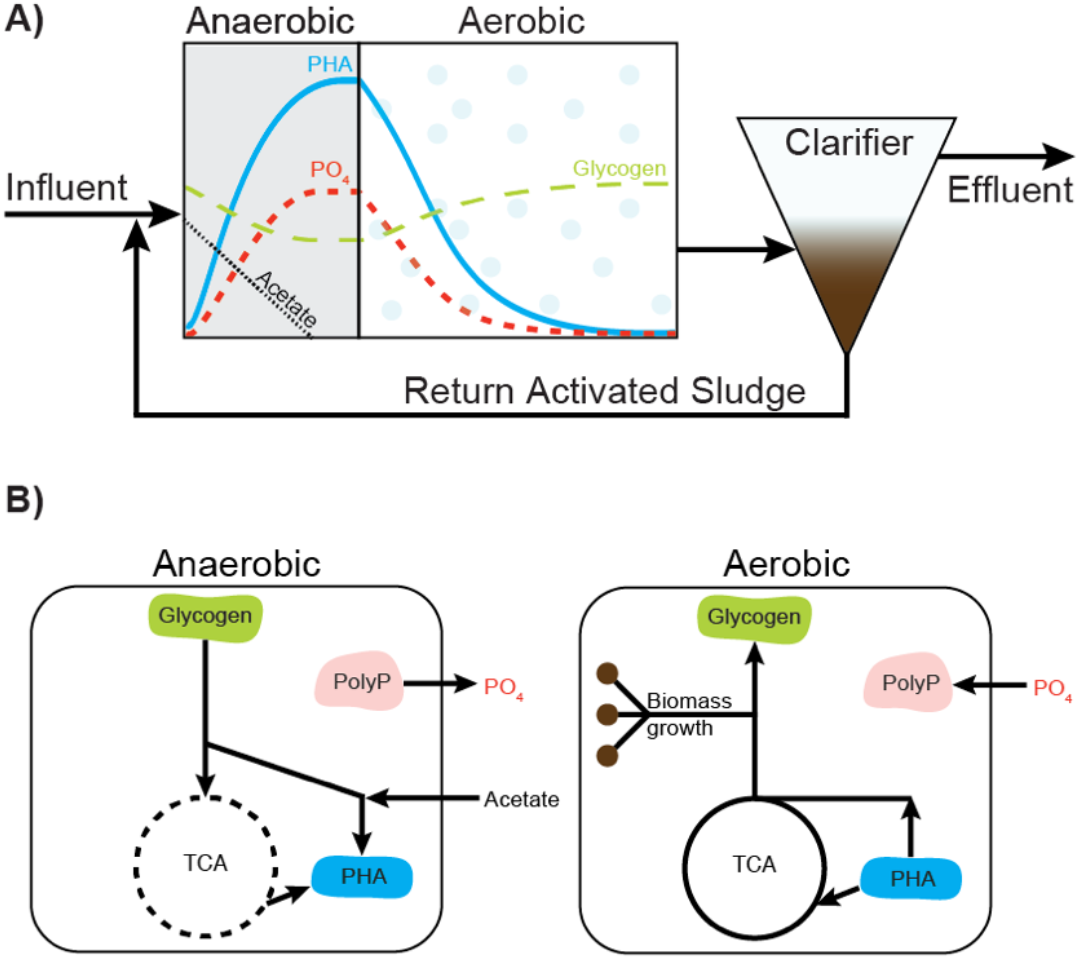
Schematic diagram of an EBPR process with nutrient/polymer profiles (A) and corresponding metabolic strategy (B). Recirculated activated sludge containing PAOs is mixed with influent wastewater in an anaerobic reactor. To compensate for the absence of an external electron acceptor, Accumulibacter use their polyphosphate and glycogen reserves to take up organic carbon sources (e.g. acetate) and accumulate them in the form of polyhydroxyalkanoates (PHA). Phosphate is released at this stage. The TCA cycle is represented by a dashed line has it can have different operating modes. When oxygen becomes available, Accumulibacter makes use of their PHA storage to grow, to replenish glycogen and polyphosphate. The re-accumulation of polyphosphate in growing biomass and subsequent purge of this biomass leads to the net removal of phosphate from wastewater. Adapted from (2, 3). PHA, polyhydroxyalkanoates; PolyP, polyphosphate; PO_4_, phosphate; TCA, tricarboxylic acid cycle.

In the past decades, a number of researchers have derived hypotheses about Accumulibacter’s (anaerobic) physiology (4, 6–8). One of the most important inconclusive discussions is the source of reducing power for the anaerobic accumulation of polyhydroxyalkanoates (PHA) from volatile fatty acids (e.g. acetate, propionate), as reviewed by Zhou *et al* (9). Most experimental approaches were adequate to study the general physiology of Accumulibacter species, however the inherent population heterogeneity has not yet been sufficiently addressed; for example, only recently researchers have reported different Accumulibacter clades showing different kinetic characteristics leading to different metabolic operations (10). Furthermore, novel meta-omics approaches were used, which allowed for Accumulibacter-targeted (culture-independent) analysis without the interference of other, non-Accumulibacter, sub-populations. A comprehensive overview of the physiological studies published on Accumulibacter can be found in the Supplementary Document S1. In one of the metatranscriptomic studies (11), Oyserman and colleagues highlighted the need for validating assumptions often made in metabolic models of Accumulibacter. In particular, they refined the discussion on reducing power sources and requirements by making a distinction between the different redox cofactors, NADH and NADPH. This distinction imposes a constraint between sources and sinks for each redox cofactor, which can only be alleviated by energy-consuming transhydrogenase(-like) mechanisms or using electrons for hydrogen production. However, in their analysis, the specificity of the involved oxidoreductases was derived from analogy with other organisms.

In Accumulibacter, reducing power is required for the conversion of acetate to 3-hydroxybutyrate (3HB), the monomer of the reserve polymer poly-3-hydroxybutyrate (PHB). The intermediate reduction step is catalyzed by the enzyme acetoacetyl-CoA reductase (AAR). In PHA accumulating bacteria such as *Cupriavidus necator* and *Zoogloea ramigera*, the preferred electron donor cofactor of the AAR is NADPH (12), which was the cofactor assumed by analogy for Accumulibacter (11). However, AARs accepting both NADH and NADPH have been reported for *Azotobacter beijerinckii* (13, 14) and *Allochromatium vinosum* (15). A brief thermodynamic feasibility analysis of the coupling between glycolysis and either NADPH- or NADH-preferring AAR is present in the Supplementary Document S2.

So far and to our knowledge, the nature of the cofactor accepted by Accumulibacter’s acetoacetyl-CoA reductase has not yet been experimentally established. In this study, we addressed this knowledge gap by measuring the NADH- and NADPH-dependent acetoacetyl-CoA reductase activities in a cell free extract from a highly enriched culture of Accumulibacter. We further extended the analysis to other key oxidoreductases involved in the anaerobic metabolism of Accumulibacter. These results were then used to update Accumulibacter’s biochemical model and as constraints in a flux balance analysis (FBA) framework. Our simulations show how the use of different pathways and storage compounds lead to metabolic flexibility. Without this flexibility, Accumulibacter’s metabolism would become very restrictive and, under unpredictable anaerobic environments, it could lead to situations were only a limited fraction of the available substrate could be consumed.

## MATERIALS AND METHODS

### Reactor operation

Two independent Accumulibacter enrichments were obtained in sequencing batch reactors (SBR). The main operational conditions are described in Table 2 and were adapted from the SBR-S as described in Welles *et al* (10).

### Cell free extracts preparation

Broth samples (10 mL) collected from the bioreactor during both anaerobic and aerobic phases were centrifuged (2500g, 10 min, 4°C) and the pellet was washed using 10 mL buffer (hereafter named Buffer 1X) containing 50 mM Tris at pH 8, 5 mM MgCl_2_, 5 mM NaCl and 5% (v/v) glycerol, and the obtained suspension was centrifuged again (2500g, 10 min, 4°C). After centrifugation, pellets were kept at −20°C for no longer than four days until further analysis. For the enzymatic assay, cellular pellets were suspended in 10 mL of Buffer 1X supplemented with 2 mM (L+D) 1,4-Dithiothreitol (DTT) and cOmplete™ mini protease inhibitor cocktail (Roche). To avoid overheating and protein denaturation, cells were kept on ice while being sonicated until the cell suspension was homogenized (i.e. no granules visible). The resulting suspension was centrifuged (15000g for at least 45 min, 4°C). The obtained supernatant, a cell free extract (CFE), was used for the enzymatic assays.

### Enzymatic assays

The reaction mixtures used for each enzymatic assay are described in Table 1. Activities were calculated using the initial rate of reduction of NAD(P)+ or oxidation of NAD(P)H, which was obtained by following the changes in the absorbance at 340 nm, at 30°C. To control the contribution of putative background reactions, reaction mixtures were monitored that contained all the components except one substrate or without addition of CFE (see “Controls” in Supplementary Document S4). The measured rates were normalized to the total protein concentration in the respective CFE. The protein concentration was measured using the Bradford dye-binding protein assay (Bio-Rad), using known concentrations of bovine serum albumin as external standards.

**Table 1.** Origin of the cell free extract (CFE), buffer, substrates, controls and references (16–24) used for each enzymatic assay. Controls used can be found in Supplementary Document S4.

| CFE | Tested enzyme | Buffer | Substrates | Reference |
| --- | --- | --- | --- | --- |
| SBR-1 | acetoacetyl-CoA reductase (AAR) | 1X, supplemented | 200 μM acetoacetyl-CoA, 200 μM NADH or 200 μM NADPH | (16) |
| SBR-2 | acetoacetyl-CoA reductase (AAR) | 1X, supplemented | 40 μM acetoacetyl-CoA, 100 μM NADH or 200 μM NADPH | (17) |
| SBR-2 | 3-hydroxybutyryl-CoA dehydrogenase (3HBDH) | 1X, supplemented | 100 μM 3-hydroxybutyryl-CoA 1000 μM NAD(P)<sup>+</sup> | (17) |
| SBR-1 | glucose-6-phosphate dehydrogenase (G6PDH) | 1X, supplemented | 2 mM glucose-6-phosphate; 2000 μM NAD<sup>+</sup> or 200 μM NADP<sup>+</sup> | (18) |
| SBR-2 | glucose-6-phosphate dehydrogenase (G6PDH) | 1X, supplemented | 4 mM glucose-6-phosphate; 400 μM NAD(P)<sup>+</sup> | |
| SBR-2 | isocitrate dehydrogenase (ICDH) | 1X, supplemented | 400 μM NAD(P)<sup>+</sup> 400 μM isocitrate | (19) |
| | malate dehydrogenase (MDH) | 1X, supplemented | 200 μM NAD(P)H 200 μM oxaloacetate | (20) |
| | malic enzyme (ME) | 1X, supplemented | 200 μM NADP 200 μM L-malate | (21) |
| | isocitrate lyase (ICL) | 40 mM HEPES, pH 7, 6 mM MgCl<sub>2</sub> | 4 mM isocitrate, 280 μM NADH 45 units lactate dehydrogenase from rabbit muscle (Roche) | (22) |
| | fumarate reductase (FR) | 50 mM HEPES, pH 7 | 15 mM fumarate 250 μM NADH | (23) |
| | α-ketoglutarate dehydrogenase (AKGDH) | 50 mM HEPES, pH 7, 1mM MgCl<sub>2</sub>, 1mM Thiamine pyrophosphate, 2.5 mM DTT | 2 mM α-ketoglutaric acid 100 μM Coenzyme A 2.5 mM NAD(P)<sup>+</sup> | (24) |

### Microbial community characterization

The microbial community present in our enrichments has been characterized by three orthogonal approaches as described below and the detailed results can be found in Supplementary Document S3.

Fluorescence *in situ* hybridization (FISH) was used to qualitatively assess the presence of Accumulibacter in the enrichment cultures. All bacteria were targeted by the EUB338 mix (general bacteria probe) (25–27). Accumulibacter clade I and II were targeted by the probes Acc-1-444 and Acc-2-444 (28), respectively. Hybridized samples were examined with a Zeiss Axioplan-2 epifluorescence microscope.

To further confirm the specific Accumulibacter clade, the presence of the gene encoding for the polyphosphate kinase I (*ppk1*) present in Accumulibacter was tested as described in (29, 30) using the primers ACCppk1-254F and ACCppk1-1376R targeting the *ppk1* gene from Accumulibacter-like bacteria (31). The *ppk1* gene sequences obtained in this study have been deposited in the GenBank database under accession numbers MH899084-MH899086.

To identify putative side-populations, 16S-rRNA gene amplicon sequencing was applied. DNA samples from cell pellets were extracted using the DNeasy UltraClean Microbial Kit (Qiagen, The Netherlands). Approximately 250 mg wet biomass was treated according to the standard protocol except an alternative lysis was implemented. This included a combination of 5 min of heat (65°C) followed by 5 min of bead-beating for cell disruption on a Mini-Beadbeater-24 (Biospec, U.S.A.). After extraction the DNA was checked for quality by gel electrophoresis and quantified using a Qubit 4 (Thermo Fisher Scientific, U.S.A.).

After quality control, samples were sent to Novogene Ltd. (Hongkong, China) for amplicon sequencing of the V3-4 region of the 16S-rRNA gene (position 341-806) on an Illumina paired-end platform. After sequencing, the raw reads were quality filtered, chimeric sequences were removed and OTUs were generated on the base of ≥ 97% identity. Subsequently, microbial community analysis was performed by Novogene using Mothur & Qiime software(V1.7.0) (32, 33). For phylogenetical determination, the most recent SSURef database from SILVA (34) was used. The microbial communities in each enrichment were compared based on the 10 most abundant OTUs with a distinctive genus (i.e. with most reads assigned to it). The 16S-rRNA gene amplicon data have been deposited in GenBank under Bioproject PRJNA490689.

### Anaerobic biochemical model of Accumulibacter

The metabolic network shown in Figure 2 is based on the ancestral genome reconstruction of Accumulibacter proposed by (2) as well as experimental observations obtained here. Additionally, an extensive literature overview supporting this biochemical model of Accumulibacter can be found in Supplementary Document S1.

**Figure 2.**
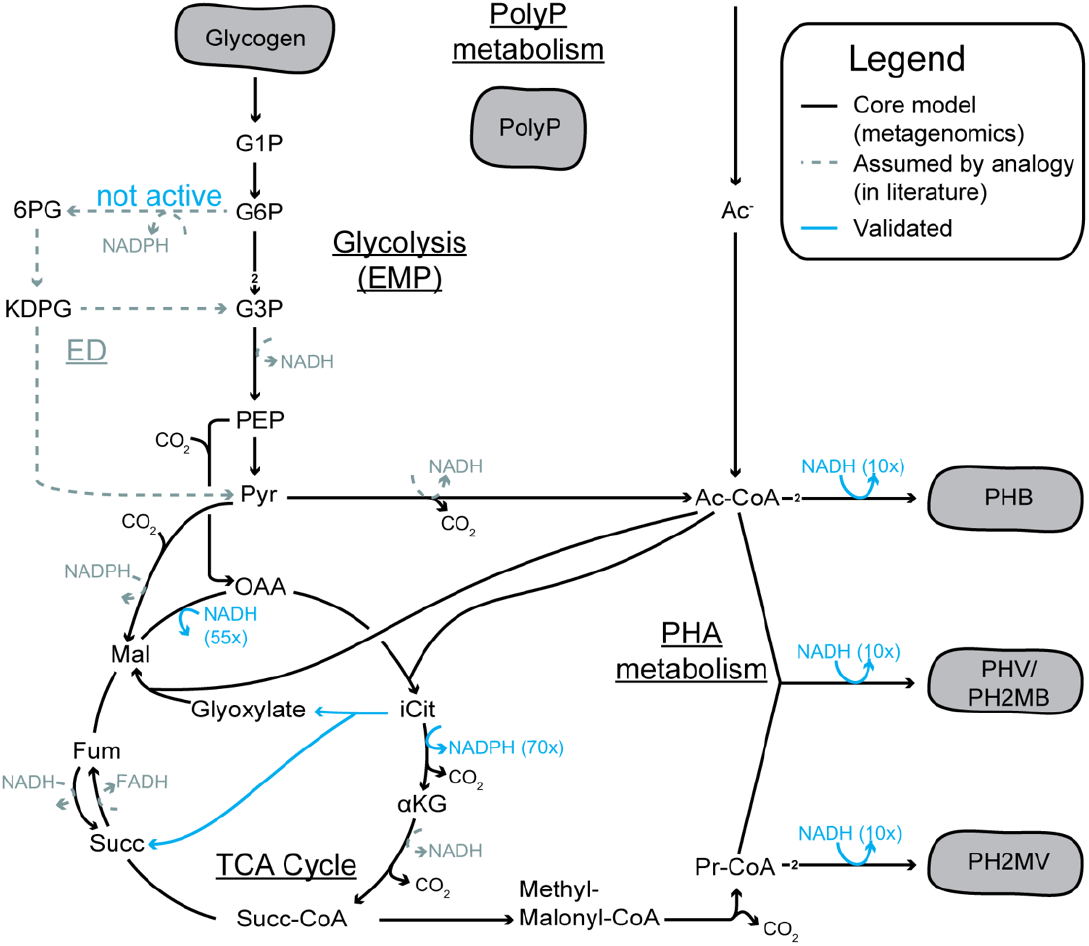
Central carbon metabolic network of Accumulibacter including redox cofactor preference of oxidoreductases assayed in this study. A simplified version of this network (Table 3) was used for the simulations. The values shown corresponds to the factor difference between the preferred cofactor and the alternative one. This metabolic network is based on the ancestral genome reconstruction done by (2). The arrows show the expected flux direction under anaerobic conditions. The untested cofactor preferences (in gray) remain as assumed by analogy with other microorganisms. Despite not being annotated in the genome, the ED pathway has been suggested as the route for glycogen degradation (40, 41). A more extensive overview of evidence for each pathway can be found in Supplementary Document S1. G1P, glucose-1-phosphate; G6P, glucose-6-phosphate; 6PG, 6-phosphogluconate; KDPG, 2-keto-3-deoxy-6-phosphogluconate; G3P, glyceraldehyde-3-phosphate; PEP, phosphoenolpyruvate; Pyr, pyruvate; OAA, oxaloacetate; Ac-CoA, Acetyl-CoA; iCit, isocitrate; αKG, α-ketoglutarate; Succ-CoA, succinyl-CoA; Succ, succinate; Fum, fumarate; Mal, malate; Pr-CoA, propionyl-CoA; Ac^−^, acetate; EMP, Embden-Meyerhof-Parnas; TCA, tricarboxylic acid cycle.

### Flux balance analysis (stoichiometric modeling)

To estimate the possible metabolic flux distributions Flux Balance Analysis (FBA) has been applied (35). The flux distribution can be obtained by optimization:

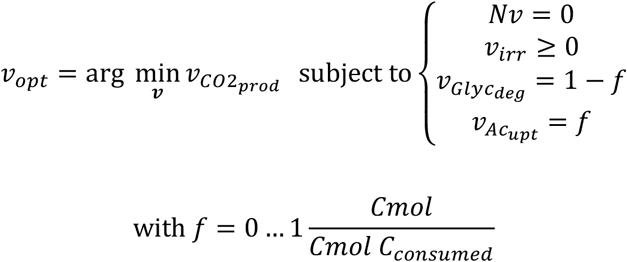

where *N* is the stoichiometry matrix containing the reactions later shown in Table 3 and *v* is the vector containing all reaction fluxes (see also Supplementary Document S6). The simulation assumes that none of the balanced intermediates is accumulating inside the cell (steady-state assumption). This is acceptable given the long simulation period compared to the turnover time of these intermediates. Inequality constraints were introduced for physiologically irreversible fluxes (i.e. these should always be positive). The consumption of acetate and glycogen is varied between only acetate (*f* = 1) to only glycogen (*f* = 0). The respective experimental data is normalized to a summed consumption of 1 Cmol (see Supplementary Document S5).

Commonly, maximization of biomass synthesis is used in FBA as an optimization objective. However, Accumulibacter only grows aerobically and on the intracellular carbon reserves (PHAs) accumulated during substrate uptake in the anaerobic period. Biomass synthesis was thus assumed proportional to the carbon stored as PHA. Consequently, maximal carbon conservation in PHA, resp. minimal CO_2_ production, is set as the cellular objective for the anaerobic phase.

To prevent adding a stoichiometric constraint due to an assumption on the proportion of the different PHA polymers possible, i.e. poly-3-hydroxybutyrate (PHB), poly-3-hydroxyvalerate (PHV), poly-3-hydroxy-2-methylbutyrate (PH2MB), or poly-3-hydroxy-2-methylvalerate (PH2MV), the respective monomer amount is introduced, i.e. the reduced precursors Ac-CoA* and Pr-CoA* such that:

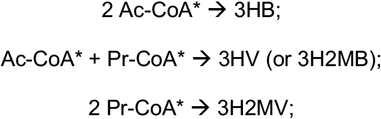

### Compilation and normalization of stoichiometric data

The results of the aforementioned simulations were compared to stoichiometric data reported in literature for several different Accumulibacter enrichments. Anaerobic-feast yields were used when available, otherwise the yields of the whole anaerobic phase were used. In case of missing compound rates (usually CO_2_, PH2MV), these were estimated using electron and carbon balancing (see Supplementary Document S5). For datasets with redundant measurements, a data reconciliation method was applied (36). For all calculations, we assumed acetate and glycogen monomers were the only substrates and CO_2_, PHB, PHV and PH2MV were the only products. In case yields were reported without the respective error, the error was calculated using error propagation. Here, the relative errors were assumed as follows: 5% for acetate and PHB measurements, and 10% for PHV, PH2MV and glycogen measurements. From the (reconciled) rates for PHB, PHV and PH2MV, the respective Ac-CoA* and Pr-CoA* rates were determined and normalized to the amount of consumed substrates, in Cmol.

## RESULTS

### Characterization of Accumulibacter enrichments

This study was carried out using two independent Accumulibacter enrichment cultures. SBR-1 contained the highest enrichment of Accumulibacter observed in our lab. However, this cultivation was operating close to a critical dilution rate which, unfortunately, resulted in washout (i.e. enrichment deterioration) after sampling. In SBR-2, conditions were adjusted to reduce the risk of washout, i.e. higher COD load and higher SRT were used.

For both cultures, anaerobic P-release per C-fed indicated that the PAO present in the sludge are likely saturated with polyphosphate and that glycogen accumulating organisms (GAO) are not present (37). Furthermore, microbial characterization by FISH showed Accumulibacter from clade I were the majority of the microorganisms across all biomass samples as it can be seen by the high overlap between the Accumulibacter-specific probe and the general Bacteria probe (micrographs available in Supplementary Document S3). The *ppk1* gene analysis further specified which subclade was dominant and the 16S-rRNA gene analysis provided information on the genus of the most abundant sub-populations next to Accumulibacter (see Table 2 and Supplementary Document S3).

**Table 2.** Process parameters and key performance indicators of the two independent enrichments of Accumulibacter.

|  | SBR-1 | SBR-2 |  |
| --- | --- | --- | --- |
| **pH** | 7.6 | 7.6 |  |
| **T** | 20°C | 20°C |  |
| **Carbon source** | 210 mg COD/L (63:37 Acetate:Propionate) | 400 mg COD/L (75:25 Acetate:Propionate) |  |
| **Phosphate load per C** | 0.1 Pmol/Cmol | 0.1 Pmol/Cmol |  |
| **Cycle time** | 6 h<br>Settling – 76 min;<br>Anaerobic – 135 min;<br>Aerobic – 135 min. | 6 h<br>Settling – 30 min;<br>Anaerobic – 112 min;<br>Aerobic – 200 min. |  |
| **HRT** | 12 h | 12 h |  |
| **SRT** | 4 days | 5.6 days |  |
| **Aerobic SRT** | 1.5 days | 3.1 days |  |
| **Anaerobic P-release per C-fed** | 0.58 Pmol/Cmol | ~0.75 Pmol/Cmol |  |
| **Anaerobic-feast length** | ~41 min | ~42 min |  |
| **FISH** | PAO Acc I | PAO Acc I |  |
| **16S rRNA gene amplicon sequencing (dominant OTUs)** | Ca_Accumulibacter<br>Rhodopseudomonas<br>Blastochloris | Start:<br>Dechloromonas<br>Ca_Accumulibacter<br>genus from<br>f__Hyphomonadaceae | End:<br>Ca_Accumulibacter<br>Chryseobacterium<br>genus from<br>o__Sphingobacteriales |
| ***ppk1* analysis** | Dominant: clade IC | Dominant: clade IA |  |

### Redox cofactor preferences of key oxidoreductases

For the simulation and interpretation of the metabolic network function it was essential to identify the cofactor specificity of relevant reactions. Enzymatic assays were performed using cell free extracts. While these assays cannot discriminate which organism has the respective activity, the performed assays still answer: 1) if an activity is not observed for the whole community, it is also not present in Accumulibacter, 2) if the cell extract shows a clear preference for NAD(H) or NADP(H) this is most probably also the preferred cofactor of Accumulibacter which is highly enriched in the culture.

The enzymatic activity assay results display a clear preference which is sufficient to annotate a specific cofactor. These are shown in blue in the metabolic network in Figure 2. Further details can be found in Supplementary Document S4. The different assay results will be presented from the sinks (PHA) to the putative sources of reducing power:

1. Redox cofactor preference (NADH or NADPH) of the PHA synthesis. The CFE from SBR-1 had a NADH preferring acetoacetyl-CoA reductase. The activity was 10x higher with NADH compared to NADPH as substrate. To further confirm this cofactor preference, the acetoacetyl-CoA reductase activity was assayed using CFEs from a second enrichment (SBR-2) and both directions of the reaction were monitored (i.e. NAD(P)H + acetoacetyl-CoA ↔ NAD(P)^+^ + 3-hydroxybutyryl-CoA).
2. Stoichiometry of glycolysis – Embden-Meyerhof-Parnas (EMP) or Entner-Doudoroff (ED)? To discriminate between these pathways, the presence of the glucose-6-phosphate dehydrogenase, the enzyme that catalyzes the first step of ED was measured. This reaction is also common to the oxidative branch of the pentose phosphate pathway (oxPP). No activity was found in the CFEs from both SBRs. The biological positive control, a CFE from *P. putida* KT2440, showed activity with both NAD^+^ and NADP^+^, as expected (38). Therefore, Accumulibacter nor the community have an active ED or oxPP pathway leaving EMP as glycolytic route.
3. Alternative NADPH sources. For many organisms oxPP is an important source of NADPH for growth (20). This activity was not found in Accumulibacter, raising the question if NADPH could be provided by other reaction like isocitrate dehydrogenase. The oxidation of isocitrate was tested using either NAD^+^ or NADP^+^. The activity with NADP^+^ was more than 70x higher, indicating that isocitrate dehydrogenase is a relevant source of NADPH in Accumulibacter.

Additionally, the activity of other oxidoreductases and anaplerotic routes was measured: i) the reduction of oxaloacetate to malate (catalyzed by malate dehydrogenase) had, at least, 55x higher activity when using NADH than NADPH. Because of the small activity observed when using NADPH, it was not possible to independently study the cofactor preference of the malic enzyme (oxidation of malate to pyruvate); ii) the glyoxylate shunt was also found to be active and to have an activity comparable to the positive biological control *E. coli* grown in acetate (well known to make use of the glyoxylate shunt (39)); and lastly, iii) fumarate reductase and α-ketoglutarate dehydrogenase were also tested but their activities were very low compared to all other enzymes tested in our study.

### Anaerobic stoichiometric model construction

Based on the defined reaction network (Figure 2 and Table 3) the balance of reducing equivalents by Accumulibacter during anaerobic acetate conversion to PHAs was studied quantitatively. Note that the network from Figure 2 was further simplified (Table 3) based on the presented evidence that PHA accumulation in Accumulibacter is NADH-consuming rather than previously assumed NADPH preference based on genome annotations. The cofactor NADH can be regenerated from other electron carriers like NADPH or ferredoxin using transhydrogenases without any energetic cost for the cell (20).

**Table 3.** Stoichiometry used for flux balance analysis (in mol basis) to simulate Accumulibacter’s <u>anaerobic</u> metabolism. Reversible reaction: ↔, irreversible reaction: →. Note that these are lumped reactions, and each may involve several enzymatic steps. A comprehensive overview of the physiological studies published on Accumulibacter supporting this stoichiometric model can be found in the Supplementary Document S1.

| Pathway | Stoichiometry | Enzyme assayed in this study |
| --- | --- | --- |
| **Glycolysis (EMP)** | $(\text{Glycogen})_1 \rightarrow 2 \text{ Pyr} + 4 \text{ electrons}$ | G6PDH |
| **Pyruvate to Acetyl-CoA** | $\text{Pyr} \rightarrow \text{Ac-CoA} + 2 \text{ electrons} + \text{CO}_2$ | |
| **Pyruvate to Oxaloacetate** | $\text{Pyr} + \text{CO}_2 \leftrightarrow \text{OAA}$ | |
| **Oxidative TCA branch** | $\text{OAA} + \text{Ac-CoA} \rightarrow \text{Succ-CoA} + 2 \text{ CO}_2 + 4 \text{ electrons}$ | ICDH<br>AKGDH |
| **Reductive TCA branch (reverse TCA)** | $\text{OAA} + 4 \text{ electrons} \rightarrow \text{Succ-CoA}$ | MDH<br>FR |
| **Glyoxylate shunt** | $2 \text{ Ac-CoA} \rightarrow \text{Succ-CoA} + 2 \text{ electrons}$ | ICL |
| **Succinate-propionate shunt** | $\text{Succ-CoA} \leftrightarrow \text{Pr-CoA} + \text{CO}_2$ | |
| **Ac-CoA\* production (precursor for 3HB, 3HV and 3HMB)** | $\text{Ac-CoA} + \text{electron} \rightarrow \text{Ac-CoA}^*$<br>(* means "+1electron") | AcAc-CoA red |
| **Pr-CoA\* production (precursor for 3H2MV, 3HV and 3HMB)** | $\text{Pr-CoA} + \text{electron} \rightarrow \text{Pr-CoA}^*$<br>(* means "+1electron") | AcAc-CoA red |

This is not the case for FADH, which would require an input of energy (ATP) to be re-oxidized using NAD^+^. Currently, there is still no experimentally validated mechanism on how Accumulibacter could re-oxidize FADH in the absence of an external electron acceptor (e.g. oxygen or nitrate). Therefore, for this simulation, we blocked this FADH producing step in the TCA cycle (i.e. Succ-CoA to Fum) and for the remaining reactions we neglected the different types of redox cofactors by simply balancing “electrons” (note that each redox cofactor carries 2 electrons and, in some publications, these are simply referred to as [H]). Since the FADH producing step in the TCA cycle is blocked, only the oxidative branch (OAA to Succ-CoA via ICDH and AKGDH) and/or the reductive branch (OAA to Succ-CoA via MDH and FR) are possible.

In contrast to electron balances, the ATP balance cannot be used as there are still too many unknowns: 1) how much ATP can be generated from efflux of ions (potassium, magnesium and phosphate) associated with polyphosphate hydrolysis; 2) how much ATP is required for acetate uptake; 3) how much is needed to upgrade redox cofactors (i.e. transfer electrons from low to high potential cofactors); 4) and how much ATP is used for cellular maintenance. Therefore, ATP is not balanced in this simulation. Nevertheless, it is important to note that the different pathways used for redox balancing will lead to different levels of ATP generation. Thus, in cases of polyphosphate limitation, the metabolism could prefer pathways with higher ATP-yield.

Experiments show that the ratio between glycogen and acetate consumption by PAOs is variable, e.g. between high/low temperature (42) or polyphosphate/glycogen availability (37, 43, 44). To reflect this variability, simulations were performed for a range of acetate and glycogen mixtures ranging from only acetate to only glycogen and results were compared to experimental data reported in literature.

For the anaerobic sequestration of acetate as PHA, acetate needs to be reduced (Ac-CoA*). The Ac-CoA* optimum in Figure 3 corresponds to the situation in which most carbon is kept inside the cell. The required reducing power can come from either glycolysis (glycogen breakdown) or glyoxylate shunt (acetate oxidation) operation. The different modes are presented in Figure 4. These modes respectively form the extreme left and right values of the optimum line in Figure 3. If glycogen degradation can supply exactly the amount of electrons needed for the uptake and conversion of acetate, then only Ac-CoA* (as PHB) is produced and there is no need to use any part of the TCA cycle (Figure 4B, and highest PHB yield in Figure 3).

**Figure 3.**
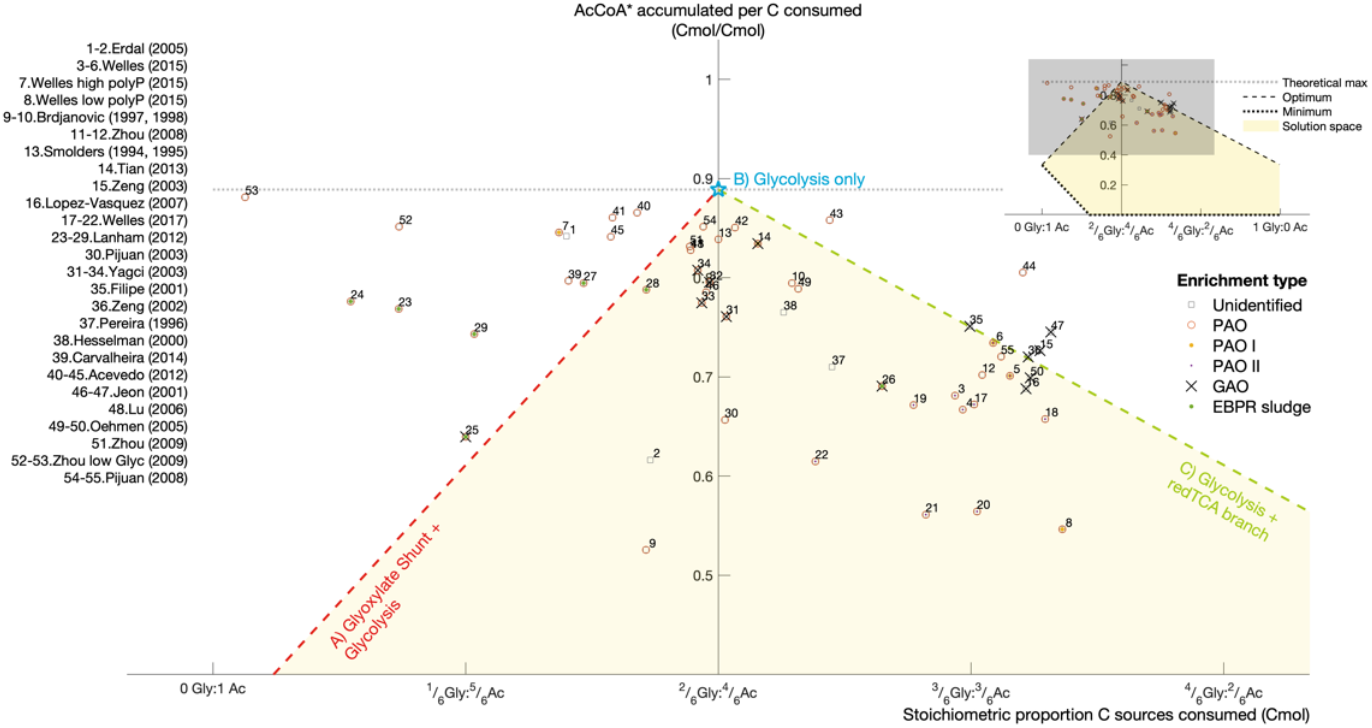
The amount of Ac-CoA* accumulated (PHB and PHV precursor) depends on the different proportion of glycogen to acetate consumed and on the available pathways. The three optimum redox balancing strategies are shown in Figure 4 ABC. From 0 to approx. 2/6 glycogen Cmol, it can be considered that bacteria are performing a polyphosphate-based metabolism (PAM, optimum line in red) and from approx. 3/6 to 1 glycogen Cmol, a glycogen-based metabolism (GAM, optimum line in green) is used. The “optimum” and “minimum” lines were obtained by minimizing and maximizing CO_2_ production, respectively. The feasible solution space (in yellow) lies in between these two cellular objectives, in which a mixture of different redox balancing strategies is used. The minimum carbon conserving strategy is the one that only uses the oxidative TCA branch. Experimental datasets were retrieved from (10, 37, 50–59, 41, 42, 44–49) and normalized to respect carbon and energy conservation principles (see Supplementary Document S5). Similar plots for Pr-CoA*, CO_2_, alternative simulations and including error bars on the experimental data points can be found in Supplementary Document S6.

**Figure 4.**
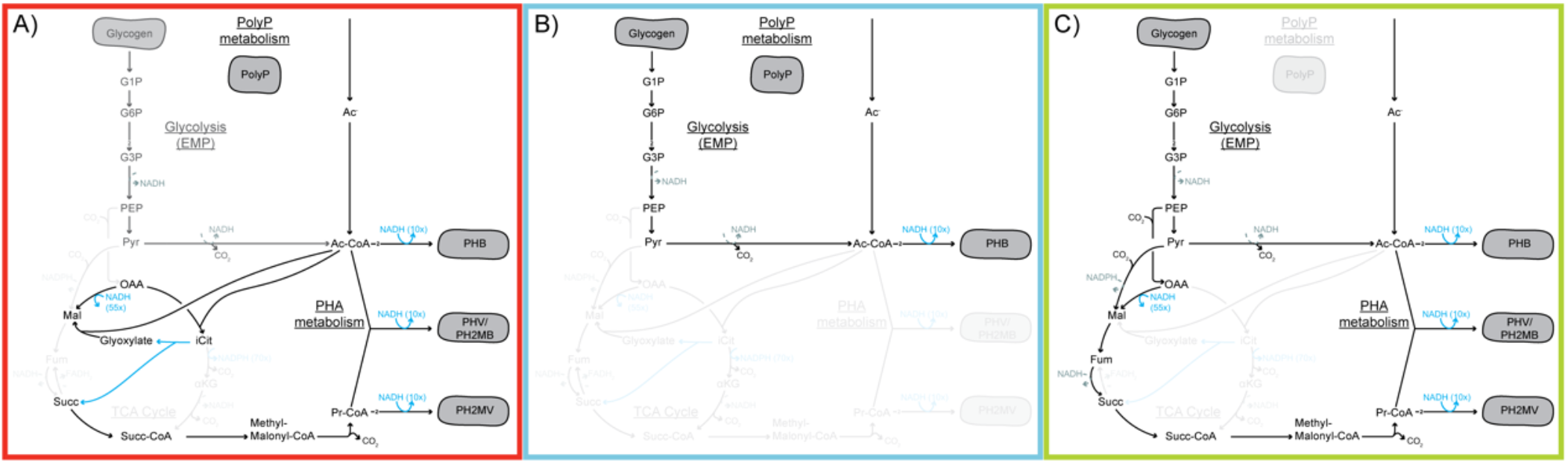
Different optimum redox balancing strategies for Accumulibacter under anaerobic conditions. A) When little glycogen is degraded and is not enough to reduce all acetate to PHB, the glyoxylate shunt as recommended by (55) is the most optimal way to provide these electrons. B) When the stoichiometric amount of glycogen is degraded for acetate reduction to PHB, there is no real need to operate any part of the TCA cycle to balance redox and only PHB is then produced, as proposed in (60). C) When more glycogen than needed for acetate reduction is degraded, the reductive TCA branch can be used as proposed by (55) for GAOs to sink electrons in more reduced PHAs (e.g. PHV and PH2MV).

The optimum can probably only be achieved with polyphosphate as ATP source. In a typical GAO-like metabolism, an excess of glycogen is degraded compared to the acetate consumed to supply all ATP needed. This leads to the generation of an excess of reducing power, which is used to produce Pr-CoA* leading to PHAs more reduced than PHB (e.g. PHV or PH2MV), using the reductive TCA branch (Figure 4C). On the other hand, in a PAO-like metabolism the amount of reducing equivalents generated by glycolysis is usually lower than required to reduce all consumed acetate. In this situation, part of the acetate is oxidized to produce reducing equivalents. Such oxidation is possible via an active glyoxylate shunt which was also found from the enzymatic assays. The simulations show this as the optimal pathway to produce PHAs when glycogen is limiting (Figure 4A), i.e. it allows for higher carbon conservation (i.e. higher PHA yield) than the “horseshoe” TCA operation.

## DISCUSSION

### Characterization of Accumulibacter enrichments

The discussion will be based on the observed high PAO enrichment of mainly *Ca*. Accumulibacter phosphatis clade I. Consequently, extrapolations of these results to Accumulibacter clade II or GAOs are speculative until further validation using respective environments for enrichments (10, 47, 61–64).

### Updated biochemical (stoichiometric) model

The presented study shows that the metabolic traits of Accumulibacter enable a flexible metabolic operation under anaerobic conditions. This flexibility is made possible by their energy and reducing power storage, enabling the observed phenotypes: fast anaerobic acetate uptake and anaerobic PHB synthesis (decoupled from growth). The flexible reducing power balancing by Accumulibacter depends on the nature of the redox cofactors used – here, enzymatic assays were performed to define the redox cofactor preferences of the main oxidoreductases in the central metabolic pathways of Accumulibacter.

The key finding from these assays is the NADH-preferring PHA accumulation in Accumulibacter. This NADH-preference allows for a direct consumption of the NADH produced in most of Accumulibacter’s reducing power sources. This also eliminates the need for NADH into NADPH conversion, which was suggested earlier by (11) using the membrane-bound transhydrogenase (PntAB) driven by proton motive force. Although there are previous reports showing NADH-driven PHB accumulation (13, 65–68), the level of NADH preference of the acetoacetyl-CoA reductase from Accumulibacter is striking. Further characterization of this enzyme was undertaken by Olavarria and colleagues (in preparation). This observation allows to re-think the role of PHAs: we hypothesize that depending on the environment where microorganisms thrive, PHA accumulation will play a role as carbon reservoir during metabolic over-flow (NADPH-driven accumulation) or as an electron reservoir during scarcity of external electron acceptors (NADH-driven accumulation). Thus, for Accumulibacter, PHA is essentially a fermentation product.

A related finding was the absence of activity of the first step of the sometimes-implicated ED glycolytic pathway. These results match those observed by (42) and in (69) for GAOs, where no NADP^+^ dependent glucose-6-phosphate dehydrogenase activity was found. The experimental findings are in line with the absence of key ED genes in the genome annotations of Accumulibacter and closest relatives *Dechloromonas aromatica* and *Azoarcus sp. EbN1* (64, 70). Furthermore, no enzymatic activity of the ED pathway with NAD^+^ as electron acceptor (38) was observed.

These findings are in contradiction with earlier ^13^C NMR studies (40, 41), which indicated that the ED pathway was more likely the route for glycolysis than EMP. Nevertheless, it has to be noted that these early studies comprise interpretations of ^13^C patterns using simplified metabolic models, with limited information on the reversibility of each reaction and potentially with reactions missing; these are common pitfalls of the ^13^C-labelling method (71). Also note that these ^13^C NMR studies were performed before the first draft genome of Accumulibacter was available (70), and only the ^13^C pattern in the different PHAs was measured and not in the metabolic intermediates of each pathway used.

While NADPH is not required anaerobically by Accumulibacter, this reducing cofactor has to be produced aerobically to drive biomass synthesis. Here, isocitrate dehydrogenase was found NADP^+^ dependent. This is consistent with the observation that most acetate consumers will use this conversion to produce NADPH for their anabolism (72) and confirms the protein annotation found in (64). Since PHA accumulation is now known to be NADH-preferring and without another sink of NADPH, the latter might accumulate anaerobically and thereby inhibit the isocitrate dehydrogenase reaction. Alternatively, a soluble transhydrogenase could convert this NADPH into NADH and allow the oxidative branch of the TCA to be operational under anaerobic conditions.

Regarding the activity of other TCA oxidoreductases and anaplerotic routes: i) the oxidation of malate using NAD^+^ was also found in the studies of (42); ii) the glyoxylate shunt was also found active as observed in assays done by (41, 42, 73) and metatranscriptomics/proteomics studies by (2, 63, 74–76), which is expected as this is the anaplerotic route that allows for microorganisms to convert C2 sources like acetate into C4 building blocks for anabolism (72); and iii) fumarate reductase and α-ketoglutarate dehydrogenase activities were very low compared to all other enzymes tested in our study and alike in the studies of (42). An α-ketoglutarate:ferredoxin oxidoreductase has been identified in Accumulibacter’s genome and proteome (64). This has not been assayed here as sufficient evidence was collected suggesting an anaerobic operation mode via the glyoxylate shunt rather than a full or partial oxidative TCA cycle.

### Flexible anaerobic metabolism – Adjustments depending on the environment and intracellular storage compounds

As (9) suggested in their review, the flexibility of Accumulibacter’s metabolism is likely the major reason why there is controversy in literature regarding how reducing power is balanced under anaerobic conditions. Long-term exposure to set conditions (e.g. pH, temperature, oxidation level of the substrate, nutrient availability, counter-ions, SRT, settling time) will select for the best strategy for those conditions, but short-term perturbations of those set conditions have shown that Accumulibacter seems to still be able to solve the redox balancing problem even if sub-optimally regarding carbon conservation (10, 43, 44, 77). Thus, based on our analysis, we observe that there is not one fixed stoichiometry, but a range of possible stoichiometries that in the end are defined by the relative proportions of each of the intervening substrates, supplied to the system or produced by a side-population (e.g. acetate, glycogen, polyphosphate, other VFAs, oxygen, nitrate, hydrogen).

The simulations showed Accumulibacter can operate in three distinct modes (and combinations in between): A) when reducing equivalents from glycolysis are limiting compared to the acetate imported (Yagci’s model (55)); B) when glycolysis supplies exactly enough reducing equivalents to convert all imported acetate into PHB (Mino’s model (60)); and lastly C) when glycolysis is producing an excess of reducing equivalents (alike Yagci’s model for GAOs (55)).

The anaerobic use of the glyoxylate shunt in a PAO-like metabolism to provide for the reducing equivalents needed for PHA accumulation is supported by the studies of (55, 73); However, as seen in Figure 3, it does not explain the higher levels of Ac-CoA* found (points above the red optimum line) when a shift to more reduced PHAs (Pr-CoA*) was expected. This indicates that there might be an alternative process that allows the cell to conserve extra carbon, in other words, accumulate more Ac-CoA* and less Pr-CoA*. It could be that 1) fully anaerobic conditions were not attained when performing the experiments, or that 2) another side population is providing electrons in the form of a more reduced organic substrate or even hydrogen gas (78) or 3) Accumulibacter has yet another, alternative way of balancing redox which has yet to be described and demonstrated.

Also for the scenario where glycolysis produces an excess of reduced cofactors as in a GAO-like metabolism, a few experimental data points fall outside the feasible solution space (points above the green optimum line in Figure 3); these could be explained in the case that Accumulibacter produces H2 to solve an excess of NADH as described by (11).

### The multilayered complexity of a physiological analysis of Accumulibacter

The experimental data gathered are likely also influenced by the intrinsic heterogeneity of Accumulibacter enrichments; clade differences, population heterogeneity or even different environment (e.g. depending on the position of the cell in a floc/granule). Therefore, experimental data points under the optimum line in Figure 3 likely represent mixtures of cells, each with a different metabolic mode (Figure 4 A, B or C). Furthermore, any kinetic limitation (i.e. pathway capacity) may also explain a sub-optimal, mixed phenotype.

Additionally, if all enzymatic machinery is available, it seems possible that the same cell might change modes depending on the environment and its intracellular storage dynamics. Particularly, availability of acetate (e.g. transition feast/famine) and/or polyphosphate storage as well as glycogen can trigger shifts in metabolic mode.

Despite the heterogeneity and noise, the experimental data outside the solution space suggests this stoichiometric model is still incomplete to represent all experimental conditions. The solution space expands for example when (1) allowing for hydrogen production in excess glycogen-to-acetate conditions or (2) the oxidation of acetate in a fully operational TCA cycle in a glycolysis-limiting scenario. With these reactions, all experimental data can be explained (see Supplementary Document S6). The first is supported by the detection of hydrogen gas produced by an Accumulibacter enrichment (11). For the latter, a full TCA cycle has been previously suggested based on ^13^C NMR observations (58) and stoichiometric analyses by (43). Additionally, a novel protein has been proposed based on metagenomic analysis that could allow for full TCA operation under anaerobic conditions (70); however, there is still no direct biochemical experimental evidence that indeed validates a full anaerobic TCA operation. Some indirect observation is presented in (79). Here, labelled propionate was used as substrate and ^13^C enrichment was found in the PHV fragments that come from acetyl-CoA. This labeling pattern can be explained by a full TCA, but this is not the only metabolic route that could explain this observation.

Alternatively, it should also be considered that, alike many (strict) anaerobes, Accumulibacter may use electron bifurcation mechanisms (80) via ferredoxin oxidoreductases to allow for alternative pathways in the central carbon metabolism.

## CONCLUSION

In this study a network-based modeling approach was used to unravel the metabolic flexibility of Accumulibacter under dynamic conditions. The approach integrates findings and hypotheses derived from previous meta-omics studies as well as different physiological datasets and biochemical assays.

The developed metabolic model demonstrates the flexibility in metabolic function and could explain previous controversy in PAO literature. The NADH dependent PHA synthesis is both an efficient link to catabolic pathways as well as key for enabling metabolic flexibility. Depending on the exact history and cultivation conditions, Accumulibacter can exhibit different metabolic phenotypes; the metabolic network can handle different combinations of carbon sources (i.e. acetate and glycogen) adjusting the use of glycolysis (EMP), different branches of the TCA cycle (incl. glyoxylate shunt) and potentially other pathways not yet considered. This poses a challenge to predictive EBPR modeling, as this implies that stoichiometry is not fixed, but variable, spanning continuously from polyphosphate to glycogen-based phenotypes.

This metabolic versatility is likely what allows Accumulibacter to be, in most situations, very close to a carbon conservation optimum, which is key to ensure their competitiveness in the dynamic environments of EBPR systems.

## ACKNOWLEDGEMENTS

These investigations were supported by the SIAM Gravitation Grant 024.002.002, the Netherlands Organization for Scientific Research (NWO). The authors would like to thank Roel van de Wijgaart, Alexandre Carnet, Hein van der Wall, Koen Verhagen, David Weissbrodt, Alex Salazar, and Thomas Abeel for their collaboration in this project. The authors would also like to thank to Sergio Tomás Martínez and Eleni Vasilakou for proofreading and Ben Oyserman for his invaluable advice on this manuscript.

