## Supplementary Document 1 for "Revealing metabolic flexibility of *Candidatus* Accumulibacter phosphatis through redox cofactor analysis and metabolic network modeling"

### S. SUPPLEMENTARY DOCUMENT

#### S1. Literature overview on Accumulibacter's biochemical model

part of the manuscript:

Running title: Metabolic flexibility of Accumulibacter

Leonor Guedes da Silva, Karel Olavarria Gamez, Joana Castro Gomes, Kasper Akkermans, Laurens Welles, Ben Abbas, Mark C.M. van Loosdrecht, Sebastian Aljoscha Wahl

Department of Biotechnology, Delft University of Technology, The Netherlands

Corresponding Author: Leonor Guedes da Silva
Address: Van der Maasweg 9, 2629HZ Delft, The Netherlands
Telephone number: +31 15 278 5307

*Table S1-1 – Overview of evidence found in literature studies on the biochemical model of Accumulibacter. Note: some studies still refer to the general term "PAO" and not Accumulibacter. Furthermore, apart from the meta-omics analysis, all other evidence is originated from enrichments of Accumulibacter and thus may contain noise from side-populations.*

| Pathway | Enzyme(s) | Type of analysis | Findings and references |
|---|---|---|---|
| **Acetate transport and activation** | Na+/H+-Acetate permease symporter | Metaproteomics | (Wilmes et al. 2008) |
| | Acetyl-CoA synthetase (ACS) | Batch test + inhibition with Ap5A | Decreased acetate uptake (Hesselmann et al. 2000) |
| | Acetyl-CoA synthetase (ACS) | Metaproteomics, metatranscriptomics | Higher protein levels than ACK (Wilmes et al. 2008), highly expressed in anaerobic acetate contact (Oyserman, Noguera, et al. 2016) |
| | Acetate kinase (ACK) | Batch test + inhibition with Ap5A | Did not affect acetate uptake (Hesselmann et al. 2000) |
| | Acetate kinase (ACK) | Metaproteomics | Lower protein levels than ACS (Wilmes et al. 2008) |
| | - | Metagenomics and metaproteomics | Present (Barr et al. 2016) |
| **Phosphate transport** | Low-affinity phosphate transporter (Pit) | Metagenomics, metatranscriptomics and metaproteomics | Found (García Martín et al. 2006; Skennerton et al. 2015); Present only in flocs (Barr et al. 2016); Two present, one high anaerobically and the other aerobically (Wilmes et al. 2008); highly expressed in high P conditions (Oyserman, Noguera, et al. 2016) |

| | | | |
|---|---|---|---|
| | High-affinity phosphate transporter (Pst) | Metagenomics, metatranscriptomics and metaproteomics | Found (Skennerton et al. 2015); Present (Barr et al. 2016; He et al. 2010; Wilmes et al. 2008); highly expressed in low P conditions (Oyserman, Noguera, et al. 2016) |
| **Polyphosphate metabolism** | PPi:PEP:AMP phosphotransferase<br>PolyP:AMP phosphotransferase | Enzymatic assay | Active (Hesselmann et al. 2000) |
| | *ppk1* (PolyP production)<br>*ppx* (PolyP hydrolysis) | Metagenomics | Present (Skennerton et al. 2015) |
| | *ppk1* | qRT-PCR | Expression increased with acetate anaerobically and in early aerobic phase, decreased notably when all phosphate was taken up aerobically (He & McMahon 2011) |
| **Full TCA cycle (anaerobically)** | - | 13C NMR, TOGA-sensor | (Pereira et al. 1996; Louie et al. 2000)<br>(Lemos et al. 2003) – propionate<br>Very high $^{13}CO_2$ fraction suggests full TCA (Zhou et al. 2009), but low $^{14}CO_2$ fraction has also been reported (Bordacs & Chiesa 1989) |
| **"Horseshoe" /Branched /Split TCA** | - | 13C NMR | (Hesselmann et al. 2000; Erdal et al. 2005) |

| | | | |
|---|---|---|---|
| **TCA cycle** | All TCA cycle enzymes<br>2-Oxoglutarate:ferredoxin oxidoreductase (OGOR)<br>Fumarate reductase (FR) | Metagenomics, metatranscriptomics and metaproteomics | Found (Oyserman, Moya, et al. 2016; Barr et al. 2016; Oyserman, Noguera, et al. 2016; Skennerton et al. 2015; García Martín et al. 2006; He & McMahon 2011; He et al. 2010; Wilmes et al. 2008; Flowers et al. 2013) |
| | Malate dehydrogenase (MDH) | Radiolabeled proteomics | Synthesized aerobically for Accumulibacter I, both anaerobic and aerobic for Accumulibacter I+II (Wexler et al. 2009) |
| | Isocitrate dehydrogenase (ICDH) | Radiolabeled proteomics | Synthesized mostly anaerobically (Wexler et al. 2009) |
| | Fumarate reductase (FR) | Batch test + inhibition with oxantel | Did not affect acetate uptake (Burow et al. 2008) |
| | Fumarate reductase (FR) | qRT-PCR | Expression increased with acetate anaerobically and in early aerobic phase (He & McMahon 2011) |
| | Succinate dehydrogenase (SDH), Other TCA enzymes except MDH | Radiolabeled proteomics | Synthesized aerobically (Wexler et al. 2009) |
| | Succinate dehydrogenase (SDH) | qRT-PCR | Expression increased anaerobically in presence of acetate (Burow et al. 2008), expression increased in early aerobic phase (He & McMahon 2011) |
| | Succinate dehydrogenase (SDH) | Batch test + inhibition with malonate | Decreased acetate uptake (Burow et al. 2008; Louie et al. 2000) |

| | | | |
|---|---|---|---|
| | Aconitase | Batch test + inhibition with fluoroacetate | Decreased acetate uptake (Burow et al. 2008; Louie et al. 2000); |
| | Aconitase | Batch test + inhibition with α-ketoglutarate | Decreased acetate uptake (Louie et al. 2000); |
| **Methylmalonyl-CoA pathway** | Methylmalonyl-CoA mutase | Metagenomics, metatranscriptomics and metaproteomics, qRT-PCR | Found (Skennerton et al. 2015; He & McMahon 2011; He et al. 2010; Wilmes et al. 2008); Induced anaerobically (He & McMahon 2011) |
| **Glyoxylate shunt** | Isocitrate lyase | Enzymatic assay | Active (Hesselmann et al. 2000) (this study)<br>Not active, but still authors suggest the shunt is active (Erdal 2002; Erdal et al. 2005) |
| | Malate synthase | Enzymatic assay | Active (Erdal 2002; Erdal et al. 2005) |
| | Isocitrate lyase | Batch test + inhibition with 3-nitropropionate and itaconate | Decreased acetate uptake (Burow et al. 2008) |
| | Aconitase<br>Isocitrate lyase | qRT-PCR | Expression increased anaerobically in presence of acetate and aerobically (Burow et al. 2008; He & McMahon 2011) |
| | Isocitrate lyase<br>Malate synthase | Metagenomics, metatranscriptomics and metaproteomics, radiolabeled proteomics | Present (Oyserman, Moya, et al. 2016; Skennerton et al. 2015; Wilmes et al. 2008; Wexler et al. 2009; He & McMahon 2011), highly expressed/synthesized in |

| | | | |
|---|---|---|---|
| | | | aerobic conditions (Oyserman, Noguera, et al. 2016; Wexler et al. 2009) |
| | Isocitrate lyase | 13C-Metaproteomics | Higher aerobically than anaerobically (Wexler et al. 2009) |
| **Glycogen utilization** | Glycogen debranching enzyme (Gde) UDP-glucose 6-dehydrogenase (ywqF) | Metagenomics and metaproteomics | (Barr et al. 2016) |
| **Glycolysis** | - | 13C NMR | (Pereira et al. 1996; Satoh et al. 1992) |
| | Glyceraldehyde-3-phosphate | Batch test + inhibition with iodoacetate | Decreased glycogen utilization (Burow et al. 2008) |
| **Glycolysis – ED** | - | 13C NMR | Most likely active (Maurer et al. 1997; Hesselmann et al. 2000) |
| | Glucose 6-P dehydrogenase | Metagenomics | Not found (García Martín et al. 2006) |
| | Glucose-6-phosphate dehydrogenase | Enzymatic assay | Not active (Erdal et al. 2005; Erdal 2002), (this study) |
| **Glycolysis - EMP** | Phosphofructokinase | Enzymatic assay | Active (Erdal 2002; Erdal et al. 2005) |
| | All steps from glycogen to pyruvate | Metagenomics, metatranscriptomics and metaproteomics, radiolabeled proteomics | Found (García Martín et al. 2006; Oyserman, Noguera, et al. 2016; Barr et al. 2016; Wilmes et al. 2008; Wexler et al. 2009; Flowers et al. 2013; He et al. 2010) |

| | | | |
|---|---|---|---|
| | Bifunctional PGK/TIM fusion protein | Metagenomics and metaproteomics | Present, more efficient than individual PGK and TIM (Barr et al. 2016) |
| **Pyruvate-“related” pathways** | Pyruvate dehydrogenase<br>Pyruvate:ferredoxin oxidoreductase (=Pyruvate synthase) | Metagenomics and metaproteomics | Present (Barr et al. 2016; Oyserman, Moya, et al. 2016) |
| | Pyruvate dehydrogenase | qRT-PCR | Expression increased in early aerobic phase (He & McMahon 2011) |
| | Pyruvate synthase | qRT-PCR | Expression increased with acetate anaerobically but decreased without acetate (He & McMahon 2011) |
| | NADP-Malic enzyme | Metagenomics and metaproteomics | Present (Barr et al. 2016) |
| | Phosphoenolpyruvate carboxykinase | Metagenomics and metatranscriptomics | Highly expressed in anaerobic acetate contact (Oyserman, Noguera, et al. 2016) |
| | Pyruvate kinase | Metagenomics, metatranscriptomics | Highly expressed in anaerobic acetate contact (Oyserman, Noguera, et al. 2016) |
| **Redox balancing enzymes** | Complex Cytochrome b/b6 (acting as reverse quinol-NAD(P) reductase | Metagenomics | Found and proposed (García Martín et al. 2006)<br>Found and pointed-out it was reported under the wrong gene-id (Flowers et al. 2013) |

| | | | |
|---|---|---|---|
| | and allowing FADH oxidation anaerobically) | | Found (Skennerton et al. 2015) |
| | | Metatranscriptomics and metaproteomics | Found (He & McMahon 2011) –> but wrong gene id targeted (Flowers et al. 2013)<br>No peptide hits (Barr et al. 2016) – possibly wrong gene id targeted? |
| | | qRT-PCR | Expression increased anaerobically in presence of acetate (Burow et al. 2008) –> but wrong gene id targeted (Flowers et al. 2013) |
| | $Na^+$-translocating NADH-quinone reductase | Metagenomics | Found (Flowers et al. 2013) |
| | RuBisCO (and other Calvin cycle enzymes) | Metagenomics, metatranscriptomics | Found (García Martín et al. 2006; Skennerton et al. 2015), not found for clade IA (Flowers et al. 2013), highly expressed in low P conditions (Oyserman, Noguera, et al. 2016) |
| | Cytoplasmic Ni-Fe hydrogen dehydrogenase<br>Membrane-bound hydrogenase | Batch test, metatranscriptomics and metagenomics | Hydrogen gas production in closed anaerobic bottle with acetate and highly expressed in anaerobic acetate contact (Oyserman, Noguera, et al. 2016) |
| | Membrane-bound transhydrogenase | Metagenomics, metatranscriptomics and metaproteomics | Present (Oyserman, Moya, et al. 2016; Barr et al. 2016), expressed (Oyserman, Noguera, et al. 2016) |

| | | | |
|---|---|---|---|
| | Soluble transhydrogenase | Metagenomics | Present (Oyserman, Moya, et al. 2016) |
| **Other** | Glutamine synthetase | Radiolabeled proteomics | Synthesized mostly anaerobically (Wexler et al. 2009) |
| | Acetyl-CoA acetyltransferase (fatty acid β-oxidation) | Radiolabeled proteomics | Synthesized mostly anaerobically (Wexler et al. 2009) |

Barr, J.J. et al., 2016. Metagenomic and metaproteomic analyses of Accumulibacter phosphatis-enriched floccular and granular biofilm. *Environmental microbiology*, 18(1), pp.273–87.

Bordacs, K. & Chiesa, S.C., 1989. Carbon Flow Patterns in Enhanced Biological Phosphorus Accumulating Activated Sludge Cultures. *Water Science and Technology*, 21(4–5), pp.387–396.

Burow, L.C., Mabbett, A.N. & Blackall, L.L., 2008. Anaerobic glyoxylate cycle activity during simultaneous utilization of glycogen and acetate in uncultured Accumulibacter enriched in enhanced biological phosphorus removal communities. *The ISME journal*, 2(10), pp.1040–51.

Erdal, Z.K., 2002. *An Investigation of the Biochemistry of Biological Phosphorus Removal Systems: Biochemistry of the Enhanced Biological Phosphorus Removal Systems, chapter III: Biochemistry of the enhanced biological phosphorus removal systems*. Virginia Polytechnic Institute and State University, Blacksburg, Virginia, USA.

Erdal, Z.K., Erdal, U.G. & Randall, C.W., 2005. Biochemistry of enhanced biological phosphorus removal and anaerobic COD stabilization. *Water Science and Technology*, 52(10–11), pp.557–567.

Flowers, J.J. et al., 2013. Comparative genomics of two “Candidatus Accumulibacter” clades performing biological phosphorus removal. *The ISME journal*, 7(12), pp.2301–14.

García Martín, H. et al., 2006. Metagenomic analysis of two enhanced biological phosphorus removal (EBPR) sludge communities. *Nature biotechnology*, 24(10), pp.1263–1269.

He, S. et al., 2010. Metatranscriptomic array analysis of 'Candidatus Accumulibacter phosphatis'-enriched enhanced biological phosphorus removal sludge. *Environmental Microbiology*, 12, pp.1205–1217.

He, S. & McMahon, K.D., 2011. "Candidatus Accumulibacter" gene expression in response to dynamic EBPR conditions. *The ISME journal*, 5, pp.329–340.

Hesselmann, R.P.X. et al., 2000. Anaerobic metabolism of bacteria performing enhanced biological phosphate removal. *Water Research*, 34(14), pp.3487–3494.

Lemos, P.C. et al., 2003. Metabolic Pathway for Propionate Utilization by Phosphorus-Accumulating Organisms in Activated Sludge: 13C Labeling and In Vivo Nuclear Magnetic Resonance. *Applied and Environmental Microbiology*, 69(1), pp.241–251.

Louie, T.M. et al., 2000. Use of metabolic inhibitors and gas chromatography/mass spectrometry to study poly-β-hydroxyalkanoates metabolism involving cryptic nutrients in enhanced biological phosphorus removal systems. *Water Research*, 34(5), pp.1507–1514.

Maurer, M. et al., 1997. Intracellular carbon flow in phosphorus accumulating organisms from activated sludge systems. *Water Research*, 31(4), pp.907–917.

Oyserman, B.O., Moya, F., et al., 2016. Ancestral genome reconstruction identifies the evolutionary basis for trait acquisition in polyphosphate accumulating bacteria. *The ISME journal*, 10(12), pp.2931–2945.

Oyserman, B.O., Noguera, D.R., et al., 2016. Metatranscriptomic insights on gene expression and regulatory controls in Candidatus Accumulibacter phosphatis. *The ISME journal*, 10(4), pp.810–22.

Pereira, H. et al., 1996. Model for carbon metabolism in biological phosphorus removal processes based on in vivo 13C-NMR labelling experiments. *Water Research*, 30(9), pp.2128–2138.

Satoh, H., Mino, T. & Matsuo, T., 1992. Uptake of organic substrates and accumulation of polyhydroxyalkanoates linked with glycolysis of intracellular carbohydrates under anaerobic conditions in the biological excess phosphate removal processes. *Water Science and Technology*, 26(5–6), pp.933–942.

Skennerton, C.T. et al., 2015. Expanding our view of genomic diversity in Candidatus Accumulibacter clades. *Environmental Microbiology*, 17(5), pp.1574–1585.

Wexler, M., Richardson, D.J. & Bond, P.L., 2009. Radiolabelled proteomics to determine differential functioning of Accumulibacter during the anaerobic and aerobic phases of a bioreactor operating for enhanced biological phosphorus removal. *Environmental Microbiology*, 11, pp.3029–3044.

Wilmes, P. et al., 2008. Community proteogenomics highlights microbial strain-variant protein expression within activated sludge performing enhanced biological phosphorus removal. *The ISME journal*, 2, pp.853–864.

Zhou, Y. et al., 2009. Involvement of the TCA cycle in the anaerobic metabolism of polyphosphate accumulating organisms (PAOs). *Water Research*, 43(5), pp.1330–1340.
