## Supplementary Document 2 for "Revealing metabolic flexibility of *Candidatus* Accumulibacter phosphatis through redox cofactor analysis and metabolic network modeling"

### S. SUPPLEMENTARY DOCUMENT

#### S2. Thermodynamics of reducing equivalents balancing in Accumulibacter

part of the manuscript:

Running title: Metabolic flexibility of Accumulibacter

Leonor Guedes da Silva, Karel Olavarria Gamez, Joana Castro Gomes, Kasper Akkermans, Laurens Welles, Ben Abbas, Mark C.M. van Loosdrecht, Sebastian Aljoscha Wahl

Department of Biotechnology, Delft University of Technology, The Netherlands

Corresponding Author: Leonor Guedes da Silva
Address: Van der Maasweg 9, 2629HZ Delft, The Netherlands
Telephone number: +31 15 278 5307

To analyze this question under debate, the conversion of acetate to 3-hydroxybutyrate (3HB), the monomer of the reserve polymer poly-3-hydroxybutyrate (PHB), can be used as a starting point (for simplification, only acetate is considered as the carbon source ($C_2H_4O_2$, 4 e-/C) and 3-hydroxybutyrate (3HB) and 3-hydroxyvalerate (3HV) as monomers for PHAs ($C_4H_8O_3$, 4.5 e-/C and $C_5H_{10}O_3$, 4.8 e-/C, respectively):

$$2 \text{ Acetate} + 2 \text{ e-} \rightarrow \text{3HB}.$$

To maintain the electron supply during acetate conversion to PHA, Accumulibacter have to harvest electrons elsewhere in their metabolism. To date, the following sources have been proposed (and thoroughly reviewed in (Zhou et al. 2009)):

a. Glycogen oxidation to acetyl-CoA and $CO_2$;

b. Acetate oxidation to $CO_2$;

c. 2 Acetate oxidation to propionyl-CoA and $CO_2$;

d. Glycogen oxidation to propionyl-CoA and $CO_2$.

Anaerobic acetate oxidation alone does not provide net energy and thus this option is only possible thanks to a special feature of Accumulibacter: energy generation from the mobilization of previously stored polyphosphate. On the other hand, glycogen oxidation does generate ATP. Glycogen oxidation is actually the strategy employed by Glycogen Accumulating Organisms (GAOs) to compete with PAOs for substrate, anaerobically. Accumulibacter also make use of glycogen, especially when phosphate is scarce in their environment and polyphosphate storage is limited (PAM/GAM switch, (Acevedo et al. 2012; Acevedo et al. 2014; Welles et al. 2015; Welles et al. 2016; Zhou et al. 2008)).

Thus far, for the oxidation of glycogen, Embden-Meyerhof- Parnas (EMP) glycolysis has been confirmed several times to be the active pathway for the oxidation of glycogen into pyruvate (for evidence, see Supplementary Document 1). In this pathway, reducing equivalents in the form of NADH are produced in the reaction catalyzed by glyceraldehyde-phosphate dehydrogenase (GAPDH):

$$\text{GAP} + \text{phosphate} + \text{NAD}^+ \Leftrightarrow \text{3-phospho-D-glyceroyl phosphate} + \text{NADH}$$

In this study we found a direct link between the reducing equivalents source (glycolysis) and sink (PHA accumulation) – NAD(H) – by unravelling the redox cofactor preference of acetoacetyl-CoA reductase (AAR):

$$\text{Acetoacetyl-CoA} + \text{NADH} \Leftrightarrow \text{3HB-CoA} + \text{CoA} + \text{NAD}^+$$

For both NADH-producing and consuming reactions to run forward, the remaining reaction species are “responsible” to keep the reactions’ Gibbs free energies ($\Delta_r G$)

negative (=feasible). Thus, we asked the question *"can GAP DH (glycolysis) and AAR (PHA formation) run simultaneously when using the same redox cofactor couple, NAD/NADH?".* In Figure S2-1 one can see the reaction catalyzed by AAR has a rather broad feasible range (gray area), since one of its products, 3HB-CoA is simultaneously removed and polymerized into the solid state. Therefore, the ratio CoA/Acetoacetyl-CoA is likely what defines the direction of this reaction. The reaction catalyzed by GAPDH has a rather narrow feasible range (yellow area). Nevertheless, when overlapped, one can see the feasible range in which both GAPDH and AAR can operate simultaneously lies well within any of the typical values for the ratio NAD/NADH found in different types of cells under different conditions. Therefore, both reactions can run simultaneously.

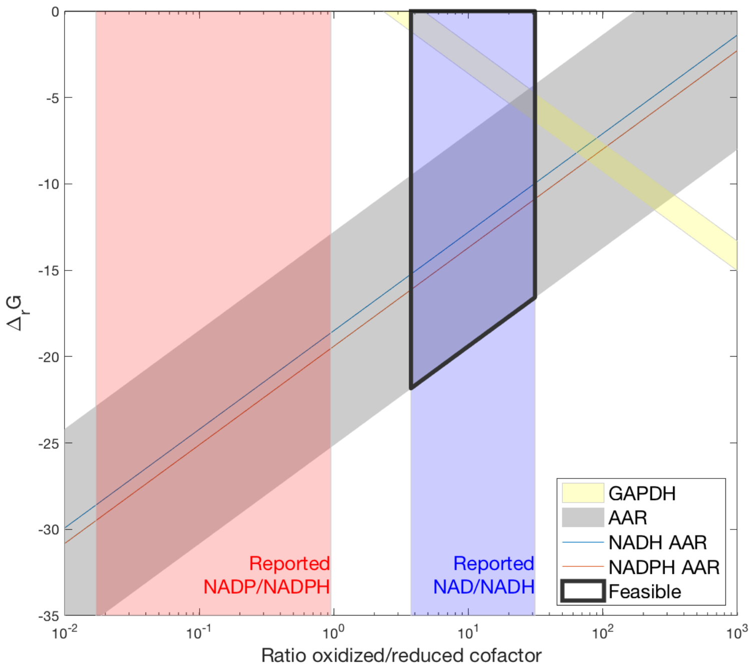


*Figure S2-1 – Thermodynamic analysis of the simultaneous operation of GAP dehydrogenase (GAPDH) and acetoacetyl-CoA reductase (AAR). Standard Gibbs free energies ($\Delta_r G^0$) were obtained from eQuilibrator* (Flamholz et al. 2012) *and the typical values for the ratios NAD(P)/NAD(P)H from* (Spaans et al. 2015)*. For the reaction catalyzed by GAP dehydrogenase, the ratio [1,3-biphosphoglycerate]/[GAP] was assumed to be 0.1 and [Pi] was varied from 50 to 100 mM based on assumptions made for other microorganisms and assuming that in PAOs, anaerobically, this value is likely higher than for a normal microorganism. For the reaction catalyzed by acetoacetyl-CoA reductase, the ratio between [3HB][CoA]/[AcAcCoA] was assumed to vary between 0.1 (more negative $\Delta_r G$) and 10 (less negative $\Delta_r G$ ).*
