## Supplementary Document 3 for "Revealing metabolic flexibility of *Candidatus* Accumulibacter phosphatis through redox cofactor analysis and metabolic network modeling"

### S. SUPPLEMENTARY DOCUMENT

#### S3. Microbial characterization

part of the manuscript:

Running title: Metabolic flexibility of Accumulibacter

Leonor Guedes da Silva, Karel Olavarria Gamez, Joana Castro Gomes, Kasper Akkermans, Laurens Welles, Ben Abbas, Mark C.M. van Loosdrecht, Sebastian Aljoscha Wahl

Department of Biotechnology, Delft University of Technology, The Netherlands

Corresponding Author: Leonor Guedes da Silva
Address: Van der Maasweg 9, 2629HZ Delft, The Netherlands
Telephone number: +31 15 278 5307

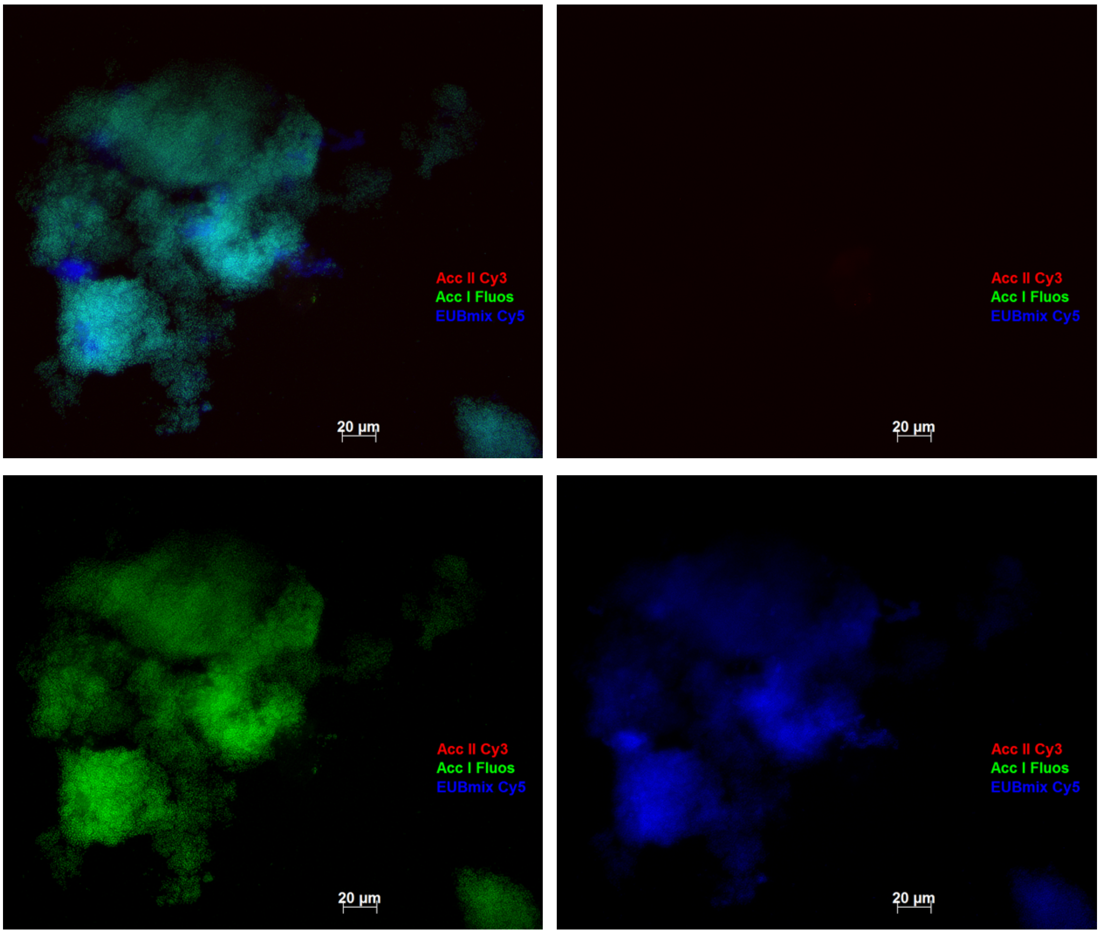


*Figure S3-1 – Standard fluorescence in situ hybridization (FISH) images from the microbial community present in SBR-1:* ***Top-left*** *All fluorescence channels merged;* ***Bottom-left*** *Cells hybridized with Accumulibacter clade I probe (Acc-1-444);* ***Top-right*** *Cells hybridized with Accumulibacter clade II probe (Acc-2-444); and* ***Bottom-right*** *Cells hybridized with the general bacteria EUB338 probes. This community was also tested with GAOmix (data not shown) and no hybridization was observed.*

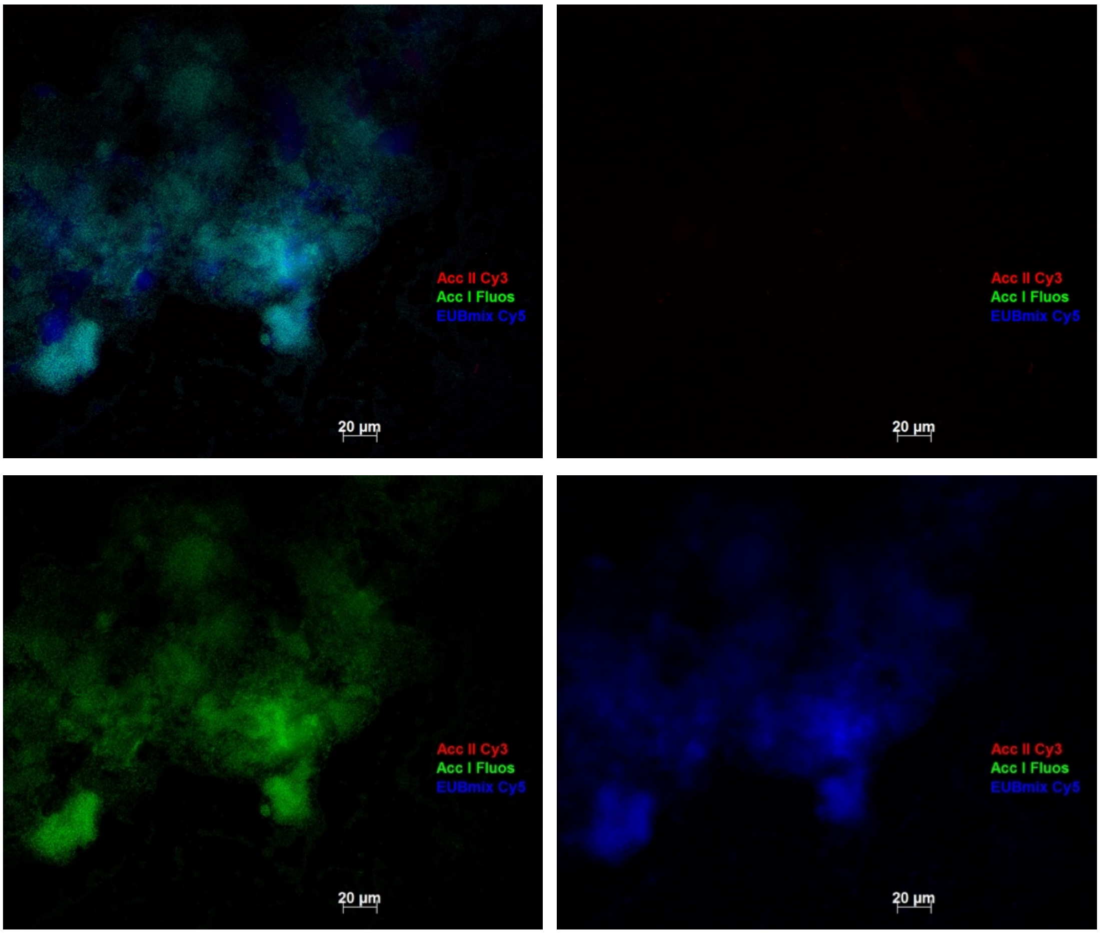


*Figure S3-2 – Standard fluorescence in situ hybridization (FISH) images from the microbial community present in SBR-2 at the start of the experiments:* ***Top-left*** *All fluorescence channels merged;* ***Bottom-left*** *Cells hybridized with Accumulibacter clade I probe (Acc-1-444);* ***Top-right*** *Cells hybridized with Accumulibacter clade II probe (Acc-2-444); and* ***Bottom-right*** *Cells hybridized with the general bacteria EUB338 probes. This community was also tested with GAOmix (data not shown) and a negligible amount of GAOs was observed.*

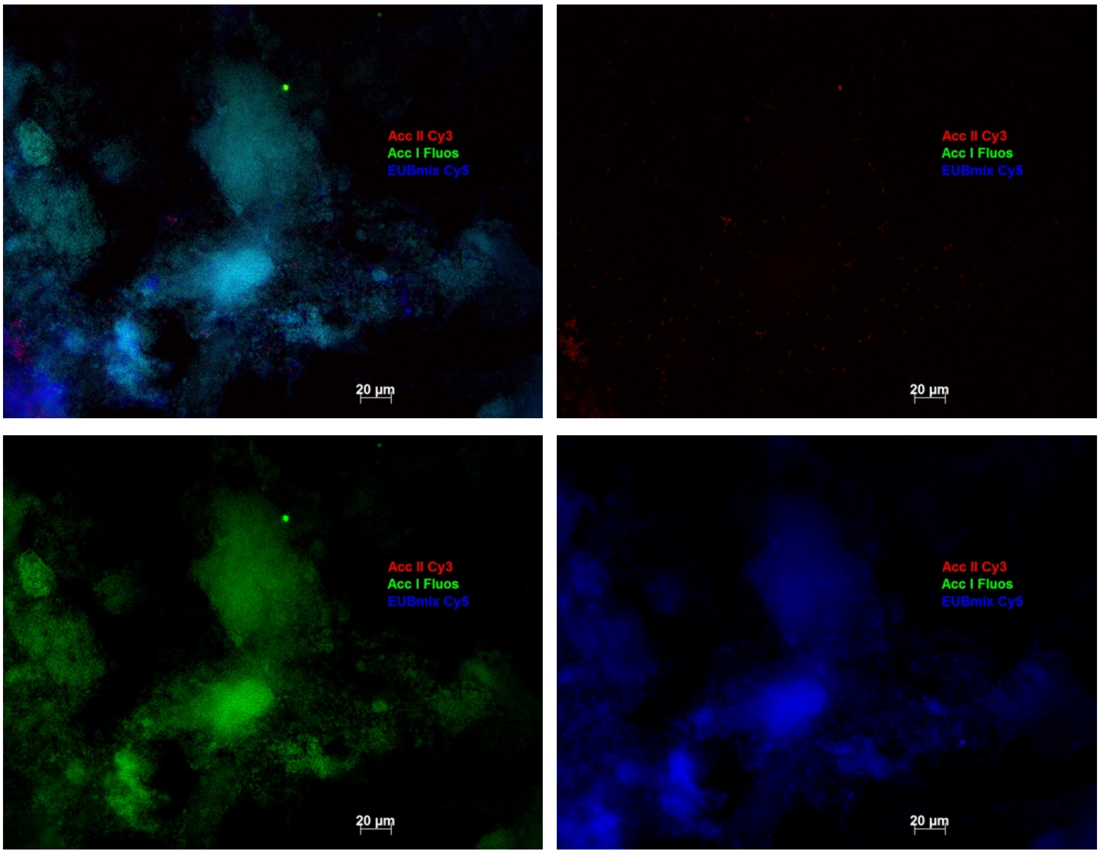


*Figure S3-3 – Standard fluorescence in situ hybridization (FISH) images from the microbial community present in SBR-2 at the end of the experiments (approx. 2 months later):* ***Top-left*** *All fluorescence channels merged;* ***Bottom-left*** *Cells hybridized with Accumulibacter clade I probe (Acc-1-444);* ***Top-right*** *Cells hybridized with Accumulibacter clade II probe (Acc-2-444); and* ***Bottom-right*** *Cells hybridized with the general bacteria EUB338 probes.*

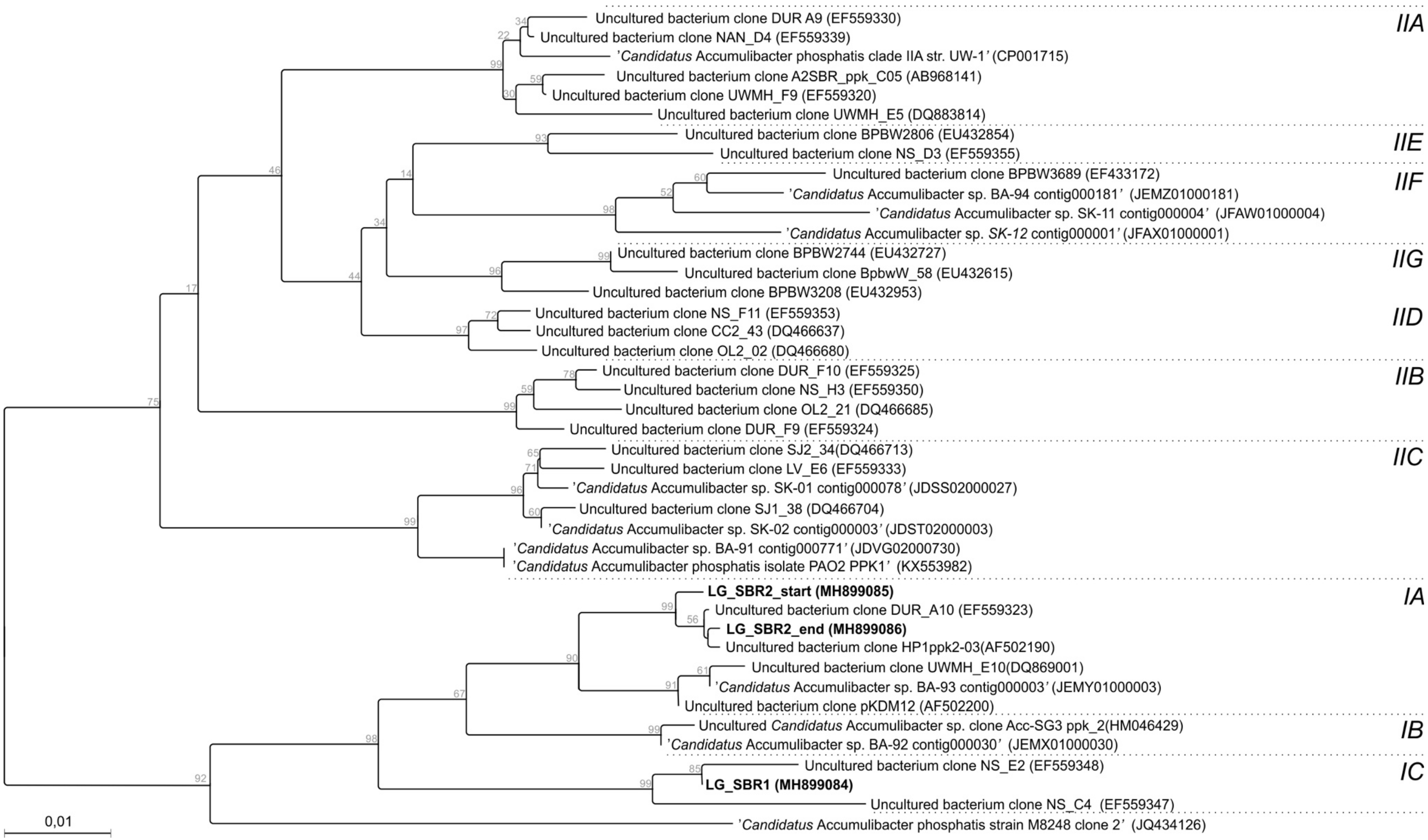


*Figure S3-4- Phylogenetic tree based on Accumulibacter's ppk gene analysis. Both SBR-1 and 2 contained Accumulibacter clade I, but subclade C and A, respectively.*

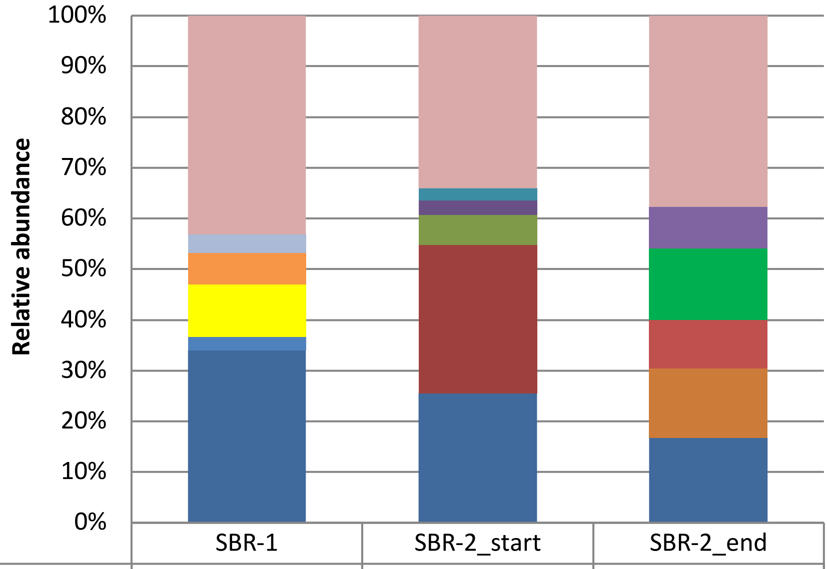


| | SBR-1 | SBR-2_start | SBR-2_end |
|---|---|---|---|
| others | 43.2 | 34.1 | 37.7 |
| Lachnotalea | 3.6 | | |
| Blastochloris | 6.2 | | |
| Rhodopseudomonas | 10.3 | | |
| Aquimonas | | | 8.3 |
| Chryseobacterium | | | 14.0 |
| Flavobacterium | | | 9.6 |
| Rhodobacter | 2.7 | | |
| genus from o__Sphingobacteriales (2) | | | 13.7 |
| Lactobacillus | | 2.4 | |
| genus from o__Sphingobacteriales | | 2.7 | |
| genus from f__Hyphomonadaceae | | 6.0 | |
| Dechloromonas | | 29.3 | |
| Candidatus_Accumulibacter | 34.1 | 25.5 | 16.7 |

*Figure S3-5 – Taxonomic distribution of microbial community based on 16s-rDNA gene copy numbers. The 16S-rRNA gene amplicon libraries have been deposited under project PRJNA490689.*
