## Supplementary Document 4 for "Revealing metabolic flexibility of *Candidatus* Accumulibacter phosphatis through redox cofactor analysis and metabolic network modeling"

### S. SUPPLEMENTARY DOCUMENT

#### S4. Enzymatic assays with cell free extracts of two independent enrichments of Accumulibacter

part of the manuscript:

Running title: Metabolic flexibility of Accumulibacter

Leonor Guedes da Silva, Karel Olavarria Gamez, Joana Castro Gomes, Kasper Akkermans, Laurens Welles, Ben Abbas, Mark C.M. van Loosdrecht, Sebastian Aljoscha Wahl

Department of Biotechnology, Delft University of Technology, The Netherlands

Corresponding Author: Leonor Guedes da Silva
Address: Van der Maasweg 9, 2629HZ Delft, The Netherlands
Telephone number: +31 15 278 5307

#### i. Controls

*Table S4-1 – Controls for each assay performed on PHA or glycolysis-related enzymes.*

| CFE | Tested enzyme | Controls |
|---|---|---|
| SBR-1 | acetoacetyl-CoA reductase | No acetoacetyl-CoA;<br>*E. coli* MG1655;<br>*E. coli* MG1655 + p-phaCAB * |
| SBR-2 | acetoacetyl-CoA reductase | No acetoacetyl-CoA;<br>Isolated/purified enzyme |
| SBR-2 | 3-hydroxybutyryl-CoA dehydrogenase | No hydroxybutyryl-CoA |
| SBR-1 | glucose-6-phosphate dehydrogenase | No glucose-6-phosphate;<br>No CFE |
| SBR-2 | glucose-6-phosphate dehydrogenase | No glucose-6-phosphate;<br>No CFE |
| SBR-2 | isocitrate dehydrogenase | No substrate;<br>No CFE |
| | malate dehydrogenase | |
| | malic enzyme | |
| | isocitrate lyase | No isocitrate,<br>no CFE;<br>*E. coli* MG1655 grown on acetate;<br>*E. coli* JW3975*[2] grown on glucose |
| | fumarate reductase | No substrate;<br>No CFE |
| | α-ketoglutarate dehydrogenase | |

* Plasmid bearing the genes *phaCAB* from *C. necator* under the control of its native promoter. These genes encode for the enzymes of the pathway leading to polyhydroxybutyrate (PHB) synthesis and it was kindly donated by Dr. J.G. Cabrera Gomez from Universidade de Sao Paulo, Brazil.
*[2] strain without isocitrate lyase from Keio collection (Baba et al. 2006):
F-, Δ(araD-araB)567, ΔlacZ4787(::rrnB-3), λ-, rph-1, Δ(rhaD-rhaB)568, ΔaceA782::kan, hsdR514

##### 1. *Purification of Accumulibacter's Acetoacetyl-CoA reductase*

A gene with high degree of similarity to *phaB* from Accumulibacter was cloned as follows. Total metagenomic DNA was isolated from aerobic granular sludge from Garmerwolde Nereda wastewater plant (Netherlands). This DNA was used as a template in a PCR reaction with primers designed according to the reference sequence of the *phaB* gene from Accumulibacter (locus CAP2UW1_3919). The primers used were PAOphaB_BamUpCrt (TCGATAGGATCCATGACGCAACGTGTTGCTTTGGTTACG) and PAOphaB_XbaDwCrt (TGTGAATCTAGATTACTGATAGTAGAGGCCACCACAG). The amplified gene was restricted with *BamH*I and cloned into the pMiniT vector using the New England Biolabs PCR Cloning Kit. The single stranded 4 base overhang resulting from the restriction was filled in by the DNA polymerase included in the kit. The plasmids from four colonies bearing the recombinant plasmid were isolated and sent for sequencing (Full 24H Sequencing, Baseclear, The Netherlands) using the primers provided with the New England Biolabs PCR Cloning Kit. Once the cloning of the isolated gene was confirmed by sequencing, it was subcloned into another vector envisioning its over-expression and purification. The vector used to overexpress and facilitate the purification of the encoded protein was pCOLA-Duet1 (Novagen). pCOLA-His6-phaB$_{PAO}$ was constructed by amplifying the chosen isolated gene using the primers phaB-PAO-nat_Bam2.FOR (TCGATAGGATCCAATGACGCAACGTGTTGCTTTGGTTACG) and phaBPAOHind.REV (CTATTAAAGCTTTTACTGATAGTAGAGGCCACCACAG) and ligating the restricted (*BamH*I and *Hind*III) amplicon in-frame with the DNA sequence encoding for the His6-tag into the MCS-1 of the pCOLA-Duet1 vector. The resulting vector pCOLA-His6-phaB$_{PAO}$ was introduced in cells of *E. coli* BL21DE3. The His-tagged proteins were purified using GE Healthcare HisTrap FF columns, following the instructions of the manufacturer. The equilibration buffer was 50 mM Tris, 100 mM NaCl, 5 mM $MgCl_2$, 5% (v/v) glycerol pH 8. For loading and washing the protein a basal imidazole concentration of 20 mM was employed and for the elution a gradient from 20 to 500 mM imidazole was used. The fractions with activities in the upper quartile were pooled, concentrated and stored with 50% glycerol until assayed for activity. The purity of the protein was assessed by SDS-PAGE.

#### ii. PHB PATHWAY

From the PHB pathway, the AcAcCoARed and the reverse reaction, 3HBCoA dehydrogenation, from a microbial community enriched with Accumulibacter were assessed, as a result the specific activities were determined as can be seen in Table S4-2.

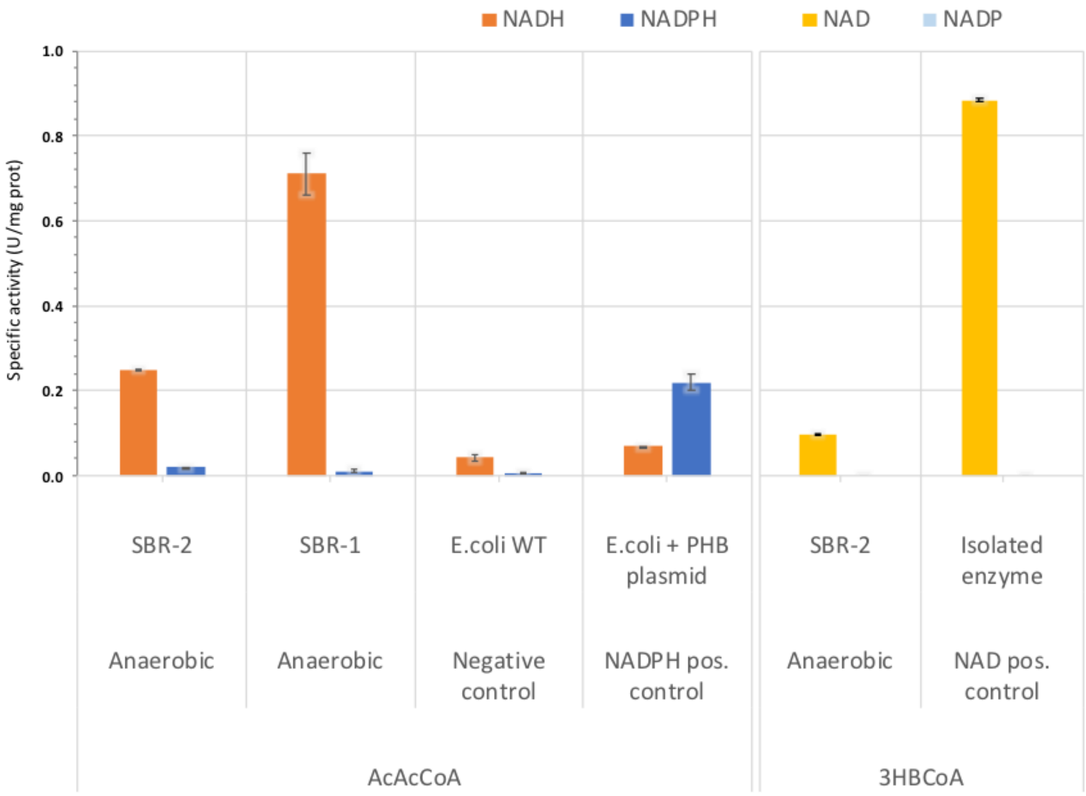

*Figure S4-1 – Accumulibacter acetoacetylCoA (AcAcCoA) reductase (step in PHA synthesis pathway) specific activity during the anaerobic phase. Note that in the assay with CFE from SBR-2 the used concentration of AcAcCoA was 5x lower, hence the lower specific activity when compared to that of SBR-1. Specific activities during the aerobic phase are comparable and thus not shown here (see Table S4-2). Controls: (1) E. coli WT was used as negative control (still, a minor activity with NAD(H) is expected since this enzyme is also involved in fatty acids synthesis and is present in E. coli.), (2) E. coli with a PHB plasmid from R. eutropha was used as NADP(H)-positive control, and (3) Purified enzyme from Accumulibacter (overexpressed and purified in E. coli as described above in Purification of Accumulibacter's Acetoacetyl-CoA reductase).*

*Table S4-2 – AcetoAcetyl-CoA reduction and 3-hydroxybutyryl-CoA dehydrogenation average specific activities (U/ mg protein) in a cell free extract from a microbial community enriched in Accumulibacter with NAD(P)H and NAD+/ NADP+, respectively, during anaerobic and aerobic phases. Standard deviations were calculated based on at least 2 technical replicates. Due to absorption of acetoacetyl-CoA at 340nm, absorbance was also measured at 310 nm for comparison but it had no effect in the conclusion regarding the cofactor specificity of this enzyme. Refer to the Materials & Methods for the used conditions in each assay. The clear cofactor preferences are marked in* ***bold****.*

| Substrate | Type CFE | CFE | Cofactor | Sp. activity NAD(P)H 340nm (U/mg prot) | SD | Sp. activity NAD(P)$^+$ 340nm (U/mg prot) | SD | Sp. activity NAD(P)H 310nm (U/mg prot) | SD |
|---|---|---|---|---|---|---|---|---|---|
| AcAcCoA | Anaerobic | SBR-2 | **NADH** | **0.249** | **0.003** | | | 0.35 | 0.01 |
| | Anaerobic | SBR-1 | **NADH** | **0.71** | **0.05** | | | | |
| | Negative control | *E.coli* WT | **NADH** | **0.042** | **0.008** | | | | |
| | NADPH pos. control | *E.coli* + PHB plasmid | NADH | 0.0691 | 0.0008 | | | | |
| 3HBCoA | Anaerobic | SBR-2 | **NAD** | | | **0.098** | **0.001** | | |
| | NAD pos. control | Isolated enzyme | **NAD** | | | **0.884** | **0.003** | | |
| AcAcCoA | Aerobic | SBR-2 | **NADH** | **0.2** | **9E-16** | | | **0.199** | **0.003** |
| | Aerobic | SBR-1 | **NADH** | **0.8** | **0.1** | | | | |
| 3HBCoA | Aerobic | SBR-2 | **NAD** | | | **0.226** | **0.009** | | |
| AcAcCoA | Anaerobic | SBR-2 | NADPH | 0.020 | 0.001 | | | 0.053 | 0.009 |
| | Anaerobic | SBR-1 | NADPH | 0.010 | 0.004 | | | | |
| | Negative control | *E.coli* WT | NADPH | 0.0060 | 0.0004 | | | | |
| | NADPH pos. control | *E.coli* + PHB plasmid | **NADPH** | **0.22** | **0.02** | | | | |
| 3HBCoA | Anaerobic | SBR-2 | NADP | | | 5E-15 | 5E-15 | | |
| | NAD pos. control | Isolated enzyme | NADP | | | 8E-14 | 2E-14 | | |
| AcAcCoA | Aerobic | SBR-2 | NADPH | 0.005 | 0.002 | | | 0.024 | 0.007 |
| | Aerobic | SBR-1 | NADPH | 0.012 | 0.002 | | | | |
| 3HBCoA | Aerobic | SBR-2 | NADP | | | 0.02 | 0.01 | | |

##### i. GLYCOLYSIS – ED pathway?

In Table S4-3, the specific activities determined with NAD and NADP for the G6PDH for the anaerobic phase, the aerobic phase and the biological controls can be found.

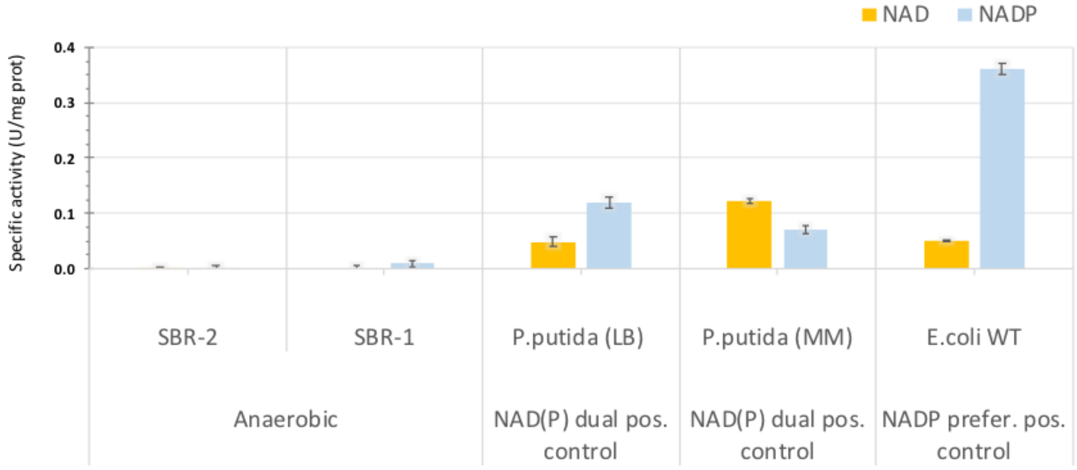

*Figure S4-2 – Glucose-6-phosphate dehydrogenase (G6PDH) specific activity (U/mg protein) in a microbial community enriched in Accumulibacter with NAD(P)+ during anaerobic and aerobic phase. Error bars represent standard deviation (4 technical replicates). CFEs from SBR-1, Pseudomonas putida (P. putida) grown in LB, P. putida grown in MM and E. coli WT were all assayed together except the SBR-2 CFE which was assayed separately later.*

*Table S4-3 – Glucose-6-phosphate dehydrogenase (G6PDH) average specific activity (U/mg protein) in a microbial community enriched in Accumulibacter with NAD(P)+ during anaerobic and aerobic phase. Standard deviations are calculated based on, at least, 2 technical replicates. CFEs from SBR-1, Pseudomonas putida (P. putida) grown in LB, P. putida grown in MM and E. coli WT were all assayed together except the SBR-2 CFE which was assayed separately later. Refer to the Materials & Methods for the used conditions in each assay.*

| Type CFE | CFE | Cofactor | Specific activity (U/mg prot) | SD |
|---|---|---|---|---|
| Anaerobic | SBR-2 | NAD | 0.002 | 0.001 |
| | SBR-1 | NAD | 0.003 | 0.005 |
| NAD(P) dual pos. control | *P. putida* (LB) | NAD | 0.049 | 0.01 |
| NAD(P) dual pos. control | *P. putida* (MM) | NAD | 0.122 | 0.004 |
| NADP prefer. pos. control | *E. coli* WT | NAD | 0.051 | 0.002 |
| Aerobic | SBR-2 | NAD | 0.07 | 0.01 |

| | | | | |
|---|---|---|---|---|
| Anaerobic | SBR-2 | NADP | 0.003 | 0.002 |
| | SBR-1 | NADP | 0.009 | 0.006 |
| NAD(P) dual pos. control | *P. putida* (LB) | NADP | 0.12 | 0.01 |
| NAD(P) dual pos. control | *P. putida* (MM) | NADP | 0.07 | 0.008 |
| NADP prefer. pos. control | *E. coli* WT | NADP | 0.36 | 0.01 |
| Aerobic | SBR-2 | NADP | 0.014 | 0.004 |

Furthermore, the GAP dehydrogenase was studied with $NAD^+$ and $NADP^+$ for both phases using a microbial community enriched in Accumulibacter and the results can be found below in Table S4-4. However, no signal was found with $NADP^+$, neither for Accumulibacter nor for the *E.coli* WT control, as expected.

*Table S4-4 – Glyceraldehyde phosphate dehydrogenase (GAPDH) average specific activity (U/mg protein) in a microbial community enriched in Accumulibacter with NAD(P)+ during anaerobic and aerobic phase. Standard deviations are calculated based on 3 technical replicates. CFEs from SBR-1 and E. coli WT were assayed. Refer to the Materials & Methods for the used conditions in each assay.*

| **Type CFE** | **CFE** | **Cofactor** | **Specific activity $NAD(P)^+$ 340nm (U/mg prot)** | **SD** |
|---|---|---|---|---|
| Anaerobic | SBR-1 | NAD | 0.32 | 0.03 |
| Aerobic | SBR-1 | NAD | 0.20 | 0.01 |
| NAD pos. control | *E.coli* WT | NAD | 0.43 | 0.01 |

##### ii. TCA CYCLE

The TCA cycle oxireductases were also assayed. The goal was mainly to determine their cofactor specificity, nevertheless, samples were taken from both anaerobic and aerobic phases to also inspect if there would be any difference in the activity of these enzymes between both phases. From the oxidative branch of the TCA cycle, isocitrate dehydrogenase (ICDH) and α-ketoglutarate dehydrogenase (aKGDH) were investigated. From the reductive (reverse) branch of the TCA, malate dehydrogenase (MDH) and fumarate reductase (FR) were tested. Finally, from the anaplerotic routes that connect the lower glycolysis to the TCA, the malic enzyme NADP-dependent (ME 2) was tested. The NAD-dependent ME could not be tested as it is not trivial to separate the contribution of NAD-dependent MDH during the assay. The resulting specific activities can be found in Table S4-5.

*Table S4-5 – Accumulibacter TCA cycle oxireductases specific activities (U/mg protein) in a microbial community enriched in Accumulibacter (only using CFEs from SBR-2) during both anaerobic and aerobic phases. No controls were used in these assays as the goal was to only assess the cofactor specificity, i.e. NAD(H) or NADP(H). Standard deviations are calculated based on, at least, 2 technical replicates. The clear cofactor preferences are marked in* ***bold****.*

| **Assayed enzyme** | **Substrate** | **Type CFE** | **Cofactor** | **Specific activity NAD(P)(H) 340nm (U/mg prot)** | **SD** |
|---|---|---|---|---|---|
| ICDH | Isocitrate | Anaerobic | NAD | 0.011 | 0.002 |
| | | Aerobic | NAD | 0.01 | 0.01 |
| | | **Anaerobic** | **NADP** | **0.80** | **0.02** |
| | | **Aerobic** | **NADP** | **1.02** | **0.05** |

| | | | | | |
|---|---|---|---|---|---|
| aKGDH | aKetoglutarate | Anaerobic | NAD | 0.006 | 0.002 |
| | | Aerobic | NAD | 0.0034 | 0.0003 |
| | | Anaerobic | NADP | 0.0007 | 0.0007 |
| | | Aerobic | NADP | 0.006 | 0.002 |
| MDH | Oxaloacetate | **Anaerobic** | **NADH** | **8.6** | **0.6** |
| | | **Aerobic** | **NADH** | **1.1** | **0.2** |
| | | Anaerobic | NADPH | 0.13 | 0.04 |
| | | Aerobic | NADPH | 0.02 | 0.04 |
| ME2 | Malate | Anaerobic | NADP | 0.096 | 0.008 |
| | | Aerobic | NADP | 0.0249 | 0.0006 |
| FR | Fumarate | Anaerobic | NADH | 0.0012 | 0.0004 |
| | | Aerobic | NADH | 0.006 | 0.004 |

##### iii. Glyoxylate shunt

Also, the presence of the glyoxylate shunt was checked by assessing the enzyme that catalyses its first step: isocitrate lyase. Here we used a coupled assay with lactate dehydrogenase which oxidizes the glyoxylate produced by the isocitrate lyase. The results can be found below in Table S4-6.

*Table S4-6 – Isocitrate lyase (ICL) specific activity (U/mg protein), measurement with an indirect method with rabbit lactate dehydrogenase (LDH), in a microbial community enriched in Accumulibacter during anaerobic and aerobic phases. Using as biological controls the ICL of E. coli WT MG1655 (strain K12) grown on MM with acetate and of E.coli JW3975 grown on MM with glucose. Standard deviations are calculated based on, at least, 2 technical replicates.*

| **Type CFE** | **CFE** | **Specific activity NAD(P)H 340nm (U/mg prot)** | **SD** |
|---|---|---|---|
| Anaerobic | SBR-2 | 0.07 | 0.01 |
| Aerobic | SBR-2 | 0.02 | 0.01 |
| Pos. control | *E.coli* WT (MM+Acetate) | 0.28 | 0.02 |
| Neg. control | *E.coli* JW3975 (MM+Glucose) | 0.01 | 0.01 |
