## Supplementary Document 5 for "Revealing metabolic flexibility of *Candidatus* Accumulibacter phosphatis through redox cofactor analysis and metabolic network modeling"

### S. SUPPLEMENTARY DOCUMENT

#### S5. Experimental data reconciliation

part of the manuscript:

Running title: Metabolic flexibility of Accumulibacter

Leonor Guedes da Silva, Karel Olavarria Gamez, Joana Castro Gomes, Kasper Akkermans, Laurens Welles, Ben Abbas, Mark C.M. van Loosdrecht, Sebastian Aljoscha Wahl

Department of Biotechnology, Delft University of Technology, The Netherlands

Corresponding Author: Leonor Guedes da Silva
Address: Van der Maasweg 9, 2629HZ Delft, The Netherlands
Telephone number: +31 15 278 5307

The experimental data collected from literature can be found in tableS5_LiteratureData_reconciled.xlsx

Data reconciliation is a method based on a weighted linear regression that gives the best estimate for measurements based on their associated error and in, this case, in such way that the overall carbon and electron balances are closed according to the model:

Substrates: Acetate and glucose (glycogen monomer)

Products: PHB, PHV, $PH_2MV$ and $CO_2$

The regression was performed using the following MATLAB script:

```
%% Experimental Data reconciliation
clear
close all
clc

%Stoichiometric matrix
                  %Ac    Gly      PHV      PHB      PH2MV      CO2
Element_Matrix = [ 1      1        1        1        1
1   ;... %Carbon
                   8/2    24/6     24/5     18/4     30/6
0/1   ]; %Electron

no_Element_balances = size(Element_Matrix,1);

%Import experimental data
literature_data_file = 'tableS5_LiteratureData_reconciled.xlsx';
[NUM,TXT] = xlsread( literature_data_file , 'All' );
Gly_Ac_exp_Cmol = NUM(:,4);                            %Cmol/CmolAc
PHV_Ac_exp_Cmol = NUM(:,5);                            %Cmol/CmolAc
PHB_Ac_exp_Cmol = NUM(:,6);                            %Cmol/CmolAc
PH2MV_Ac_exp_Cmol = NUM(:,7);                            %Cmol/CmolAc
Ac_Ac_exp_Cmol = NUM(:,3);       %Cmol/CmolAc

%Assume std dev
elemental_std = 1e-5*ones(no_Element_balances,1); %"very accurate"
- conservation of C and electrons
Ac_Ac_std  = NUM(:,8);
Gly_Ac_std = NUM(:,9);
PHV_Ac_std = NUM(:,10);
PHB_Ac_std = NUM(:,11);
PH2MV_Ac_std = NUM(:,12);

q_measured = [-Ac_Ac_exp_Cmol -Gly_Ac_exp_Cmol PHV_Ac_exp_Cmol
PHB_Ac_exp_Cmol PH2MV_Ac_exp_Cmol]';
q_std_measured = [Ac_Ac_std Gly_Ac_std PHV_Ac_std PHB_Ac_std
PH2MV_Ac_std]';
q_reconciled = [];
std_q = [];

for i=1:size(q_measured,2)

    A = [Element_Matrix ;
eye(size(q_measured(:,i),1),size(Element_Matrix,2))];
    b = [zeros(no_Element_balances,1);q_measured(:,i)];
    W = diag([ 1 ./ elemental_std ; 1 ./
q_std_measured(:,i)]); %Linear regression
```

```
    % linear regression
    q_reconciled(:,i) = inv(A' * W^2 * A) * A' * W^2 * b;

    % calculate accuracy
    A = inv( (A' * W^2 * A) ) * A' * W^2;
    cov_q = A * W^-2 * A';
    std_q(:,i) = diag( cov_q ).^0.5;

end

normalized_q_reconciled = q_reconciled ./ (-q_reconciled(1,:))
normalized_std_q = abs(normalized_q_reconciled).*
((std_q./q_reconciled).^2 +
(std_q(1,:)./q_reconciled(1,:)).^2).^0.5;

%% Write results in excel
text = {'Ac reconciled (Cmol)',...
        'Gly reconciled (Cmol)',...
        'PHV reconciled (Cmol)',...
        'PHB reconciled (Cmol)',...
        'PH2MV reconciled (Cmol)',...
        'CO2 reconciled (Cmol)',...
        'Ac std (Cmol)',...
        'Gly std (Cmol)',...
        'PHV std (Cmol)',...
        'PHB std (Cmol)',...
        'PH2MV std (Cmol)',...
        'CO2 std (Cmol)'};

xlwrite(literature_data_file,text, 'All', 'R1');

num = [ -q_reconciled(1:2,:)', ...
         q_reconciled(3:6,:)', ...
         std_q'];

xlwrite(literature_data_file, num, 'All', 'R2');

%% Print fluxes in Command Window in a nice table format
Acetate     = round(q_measured(1,:)',2,'significant');
Glycogen    = round(q_measured(2,:)',2,'significant');
PHV         = round(q_measured(3,:)',2,'significant');
PHB         = round(q_measured(4,:)',2,'significant');
PH2MV       = round(q_measured(5,:)',2,'significant');


Reconciled_Acetate     =
round(normalized_q_reconciled(1,:)',2,'significant');
Reconciled_Glycogen    =
round(normalized_q_reconciled(2,:)',2,'significant');
Reconciled_PHV         =
round(normalized_q_reconciled(3,:)',2,'significant');
Reconciled_PHB         =
round(normalized_q_reconciled(4,:)',2,'significant');
Reconciled_PH2MV       =
round(normalized_q_reconciled(5,:)',2,'significant');
Reconciled_CO2         =
round(normalized_q_reconciled(6,:)',2,'significant');

T = table(Glycogen, Reconciled_Glycogen,...
    PHV     , Reconciled_PHV, ...
    PHB     , Reconciled_PHB, ...
```

```
    PH2MV   , Reconciled_PH2MV, ...
    Reconciled_CO2);
disp(T)


%% Plot reconciled versus measured
figure(1)
plot(q_measured',normalized_q_reconciled(1:5,:)','*')
xlim([ min(min([q_measured';normalized_q_reconciled(1:5,:)']))
max(max([q_measured';normalized_q_reconciled(1:5,:)'])) ])
ylim([ min(min([q_measured';normalized_q_reconciled(1:5,:)']))
max(max([q_measured';normalized_q_reconciled(1:5,:)'])) ])
axis square
hold on
plot(xlim,ylim,'k-')
xlabel('Measured'), ylabel('Reconciled')
legend('Acetate', 'Glycogen', 'PHV',
'PHB','PH2MV','Location','SouthEast')
ax = gca; ax.YAxisLocation = 'origin'; ax.XAxisLocation = 'origin';
ax.FontSize = 14;
```
