## Supplementary Document 6 for "Revealing metabolic flexibility of *Candidatus* Accumulibacter phosphatis through redox cofactor analysis and metabolic network modeling"

### S. SUPPLEMENTARY DOCUMENT

#### S6. Flux balance analysis

part of the manuscript:

Running title: Metabolic flexibility of Accumulibacter

Leonor Guedes da Silva, Karel Olavarria Gamez, Joana Castro Gomes, Kasper Akkermans, Laurens Welles, Ben Abbas, Mark C.M. van Loosdrecht, Sebastian Aljoscha Wahl

Department of Biotechnology, Delft University of Technology, The Netherlands

Corresponding Author: Leonor Guedes da Silva
Address: Van der Maasweg 9, 2629HZ Delft, The Netherlands
Telephone number: +31 15 278 5307

##### i. Stoichiometric matrix

One of the inputs required for a flux balance analysis is a stoichiometric matrix (Table S-1) and Stoichiometry_FBA.xlsx.

*Table S-1 – Stoichiometric matrix used for the flux balance analysis to generate the "PHA optimum" and "minimum" curves. Negative numbers represent consumption, positive -> production.*

| | Glyc deg | Ac upt | CO2 prod | Glycolysis | PDH | PEPC | oxTCA | redTCA | TCAGOX | SucPr | ATP net | AcCoA prod | PrCoA prod |
|---|---|---|---|---|---|---|---|---|---|---|---|---|---|
| G6P | 1 | 0 | 0 | -1 | 0 | 0 | 0 | 0 | 0 | 0 | 0 | 0 | 0 |
| PYR | 0 | 0 | 0 | 2 | -1 | -1 | 0 | 0 | 0 | 0 | 0 | 0 | 0 |
| electrons | 0 | 0 | 0 | 4 | 2 | 0 | 4 | -4 | 2 | 0 | 0 | -1 | -1 |
| CO2 | 0 | 0 | -1 | 0 | 1 | -1 | 2 | 0 | 0 | 1 | 0 | 0 | 0 |
| AcCoA | 0 | 1 | 0 | 0 | 1 | 0 | -1 | 0 | -2 | 0 | 0 | -1 | 0 |
| OXA | 0 | 0 | 0 | 0 | 0 | 1 | -1 | -1 | 0 | 0 | 0 | 0 | 0 |
| SucCoA | 0 | 0 | 0 | 0 | 0 | 0 | 1 | 1 | 1 | -1 | 0 | 0 | 0 |
| PrCoA | 0 | 0 | 0 | 0 | 0 | 0 | 0 | 0 | 0 | 1 | 0 | 0 | -1 |
| ATP | 0 | 0 | 0 | 3 | 0 | -1 | 0 | -1 | 0 | 0 | -1 | 0 | 0 |
| rev | 0 | 0 | 0 | 0 | 0 | 1 | 0 | 0 | 0 | 1 | 1 | 0 | 0 |

The stoichiometry was checked for conserved moieties, blocked reactions and non-identifiable reactions that form internal cycles; No conserved moieties nor blocked reactions were found and the internal cycles found did not respect the reversibility constraints. After analysis, the network contained 9 independently balanced intermediates, 13 (lumped) reactions and, thus 4 degrees of freedom. ATP was included in the analysis, but was not balanced (i.e. not a constrain).

##### ii. MATLAB code

The MATLAB code for these simulations is available upon request to the authors.

##### iii. Pr-CoA* and $CO_2$

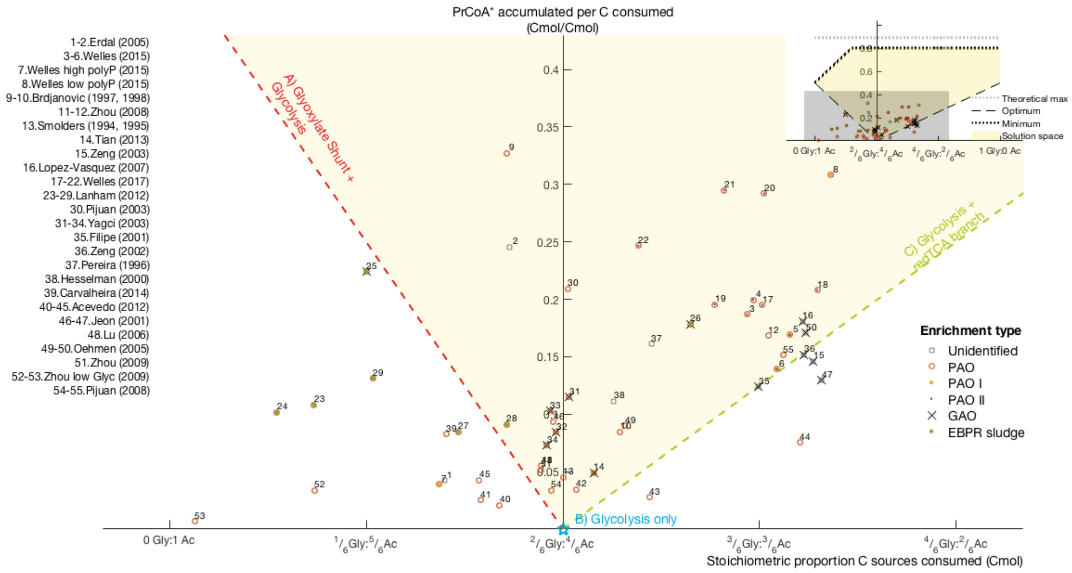


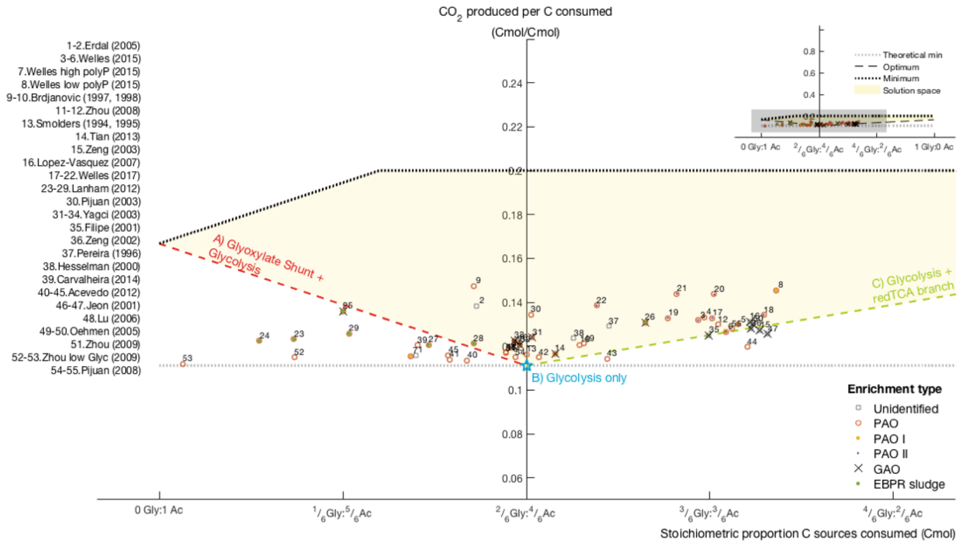


*Figure S6-1 – The amounts of Pr-CoA* accumulated and $CO_2$ released depend on the different proportion of glycogen to acetate consumed and on the available pathways.*

##### iv. Ac-CoA*, Pr-CoA* and $CO_2$ with errorbars for exp. datapoints

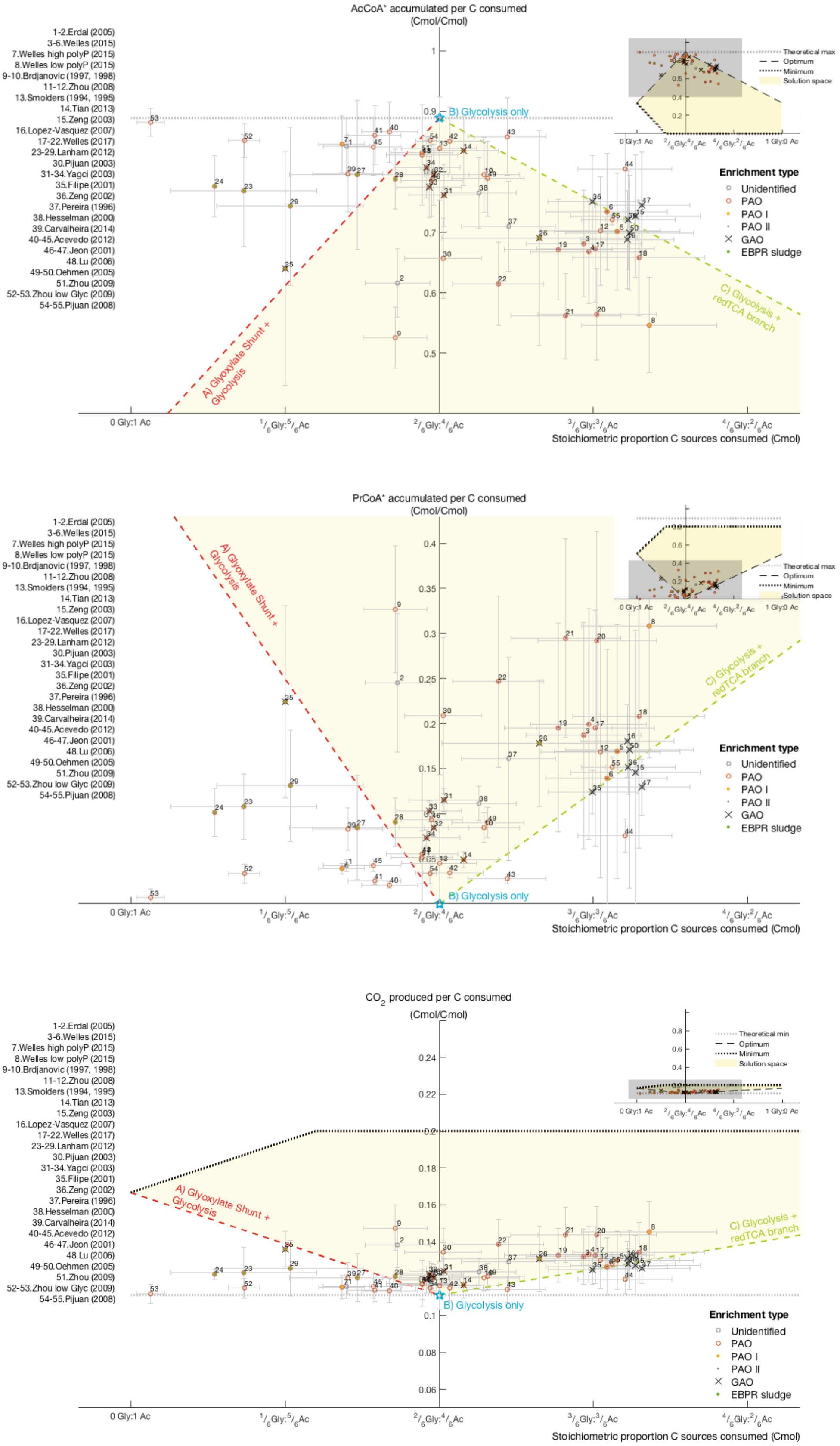


*Figure S6-2 – The error bars are propagated error based on published error when available or assumed (relative errors of 5% for acetate and PHB measurements and 10% for PHV, $PH_2MV$ and glycogen measurements).*

##### v. Electron flux distributions

Optimum scenario:

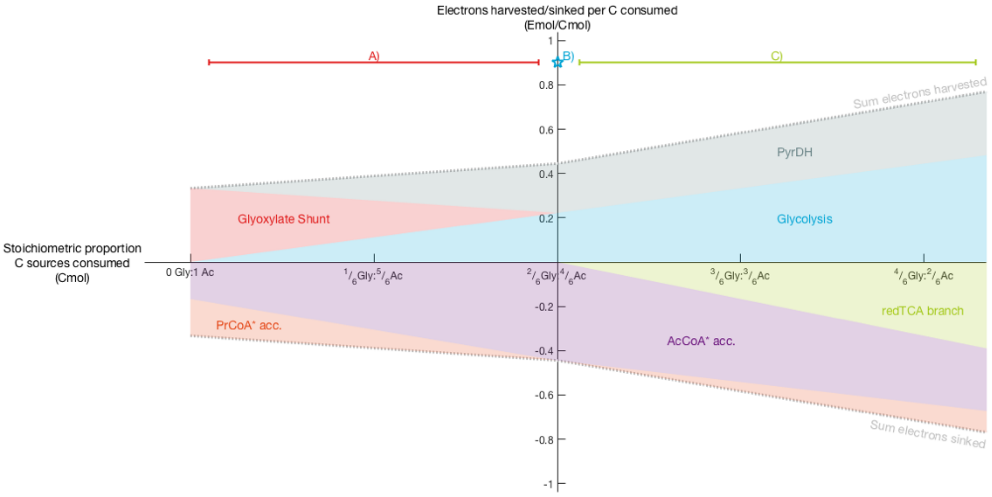


Minimum scenario:

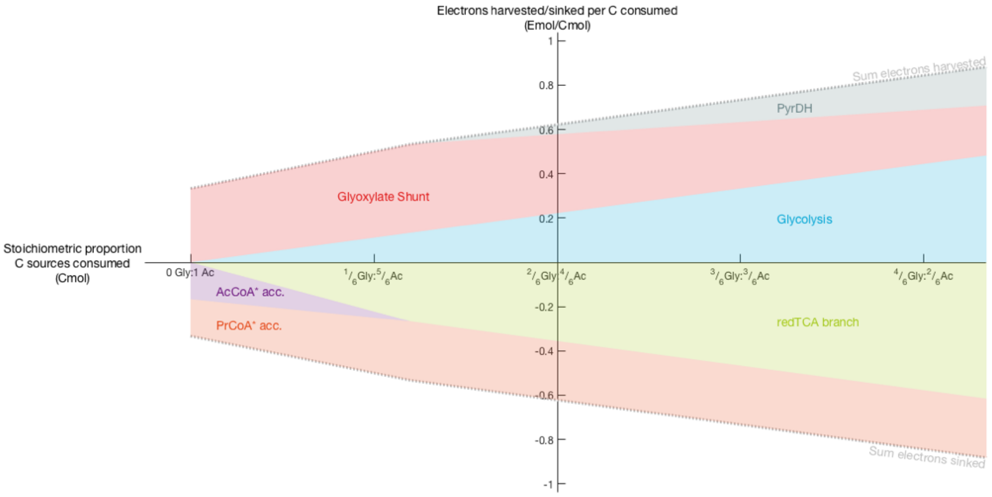


*Figure S6-3 – Electron fluxes through the different sources and sinks depending on the proportion of glycogen to acetate consumed. Values are normalized to C consumed.* ***Optimum scenario*** *– highest carbon conservation;* ***Minimum scenario*** *– lowest carbon conservation.*

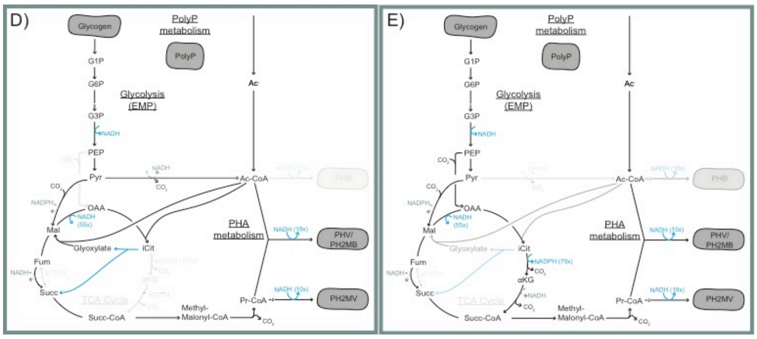


*Figure S6-4 –Lowest carbon conservation (minimum) redox balancing strategy. From another viewpoint, this is the strategy to harvest and sink the biggest amount of electrons; it makes use of both possible sources simultaneously: glyoxylate shunt and glycolysis. D) The simultaneous operation of these pathways allows for the biggest release of $CO_2$ followed closely by E) the strategy using the "horseshoe" TCA operation.*

##### vi. Comparison to Yagci (2003) PAO and GAO models

PAO model (Yagci et al. 2003):

$(6 + 3f_{GLX})$ Acetate + Glycogen -> 4 PHB + $f_{GLX}$ PHV + $(2 + f_{GLX})$ $CO_2$ with $f_{GLX} = 0{:}1000$

GAO model (Yagci et al. 2003):

Acetate + $(1/3 + 2/3\ \alpha_{GAO})$ Glycogen -> 2/3 PHB + $(1/6 + 2/3\ \alpha_{GAO})$ PHV + + $(1/2 + 2/3\ \alpha_{GAO})CO_2$ with $\alpha_{GAO} = 0{:}1000$

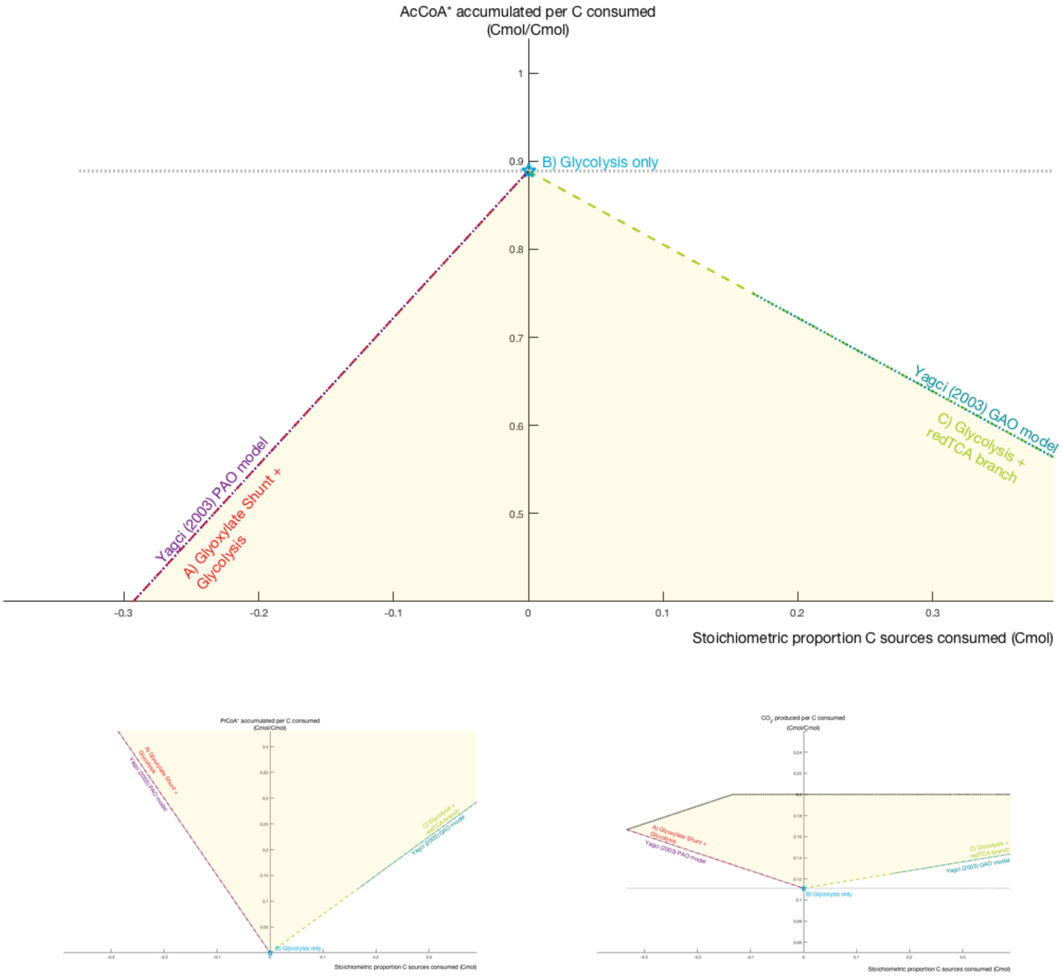


*Figure S6-5 – Our simulations and the Yagci (2003) models of PAO and GAO overlap. According to our observation, the PAO model should be extended to include the C) option until Yagci's GAO model and be renamed to PAM. The segment of C) that overlaps with Yagci's GAO model should be renamed to GAM.*

##### vii. Alternative scenarios: $H_2$ production?

The production of hydrogen gas has been observed in the studies of (Oyserman, Noguera, et al. 2016). This is a possible way of solving excess reducing equivalents. To simulate this, we added an electron sink (H2Prod) to the stoichiometric matrix, as follows:

H2Prod: 2 electrons ->

The optimization for maximum carbon conservation yielded the same as before, however now the minimum is shifted to allow more $CO_2$ release. As this would not explain any of the experimental datapoints outside the solution space, we tested the maximum Ac-CoA* (=PHB) accumulation as optimization goal. This could be the case when there is, for example, a kinetic limitation on the redTCA branch preventing excess electrons of being absorbed into Pr-CoA. The results are shown in Figure S6-6.

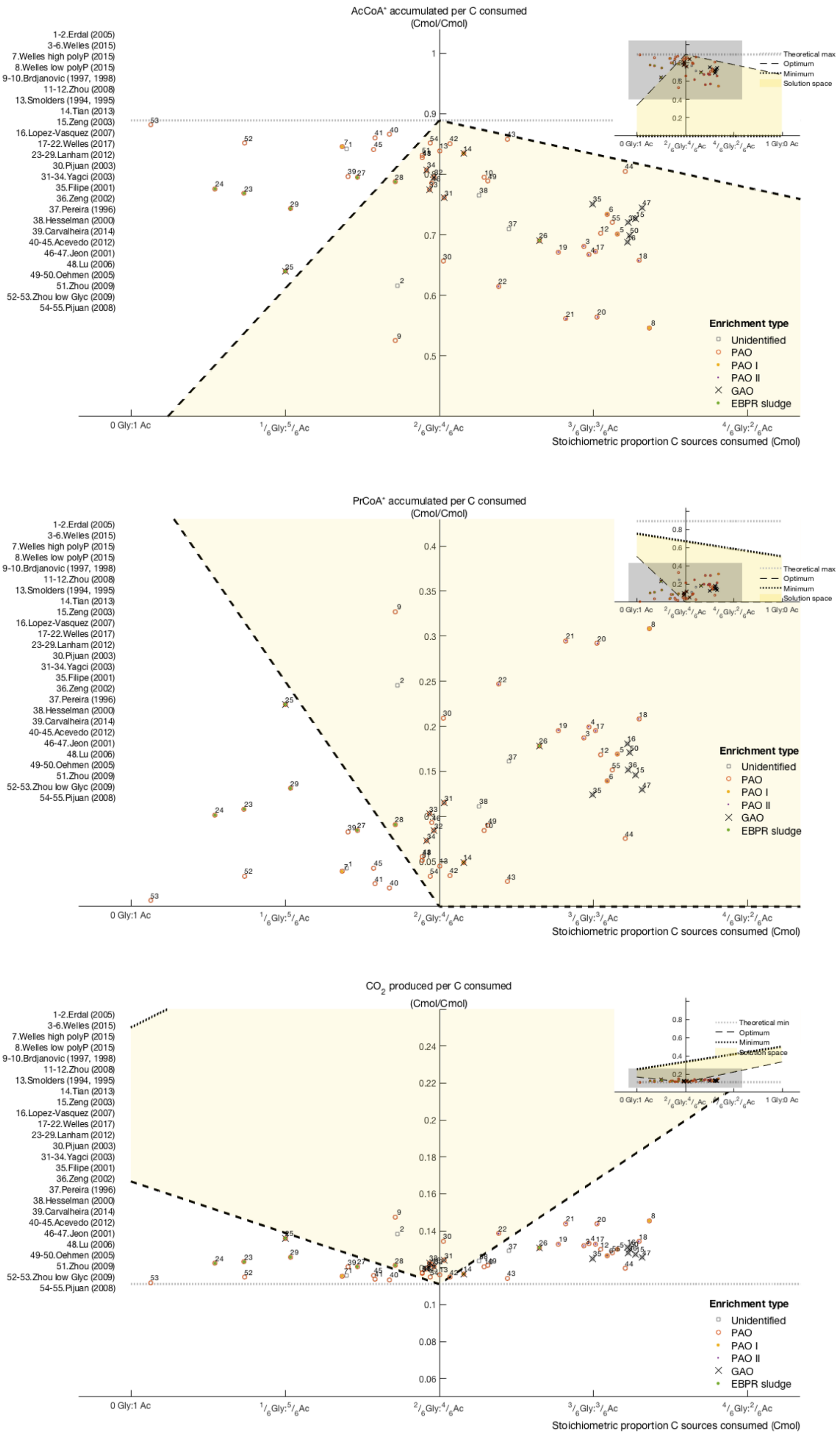


*Figure S6-6 – The $H_2$ production allows to explain the amounts of Ac-CoA* and Pr-CoA* found experimentally under the conditions of excess glycogen over acetate consumed, however it leads to higher $CO_2$ production which does not match experimental data.*

The electron flux distributions can be found below.

Optimum scenario:

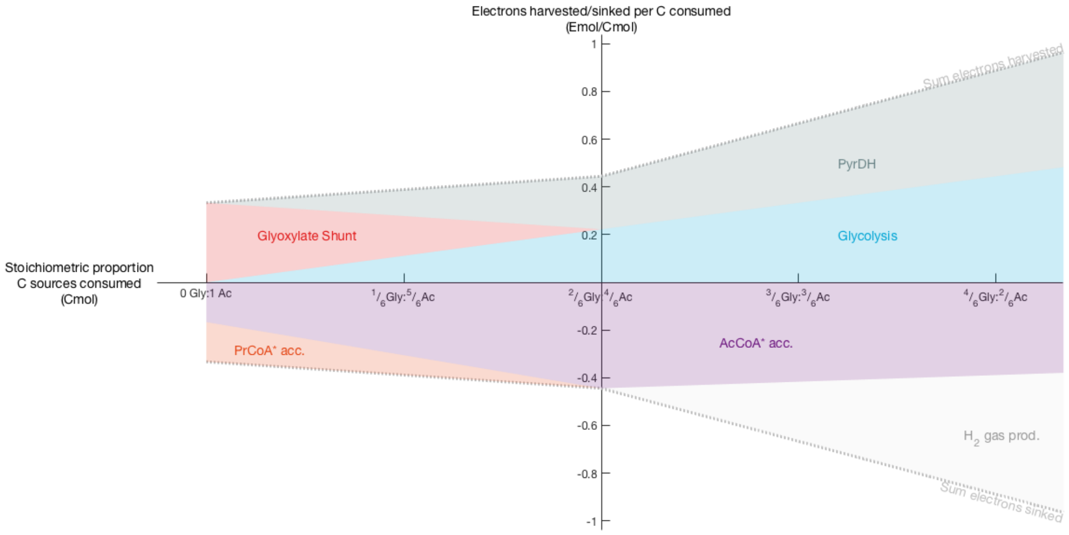


Minimum scenario:

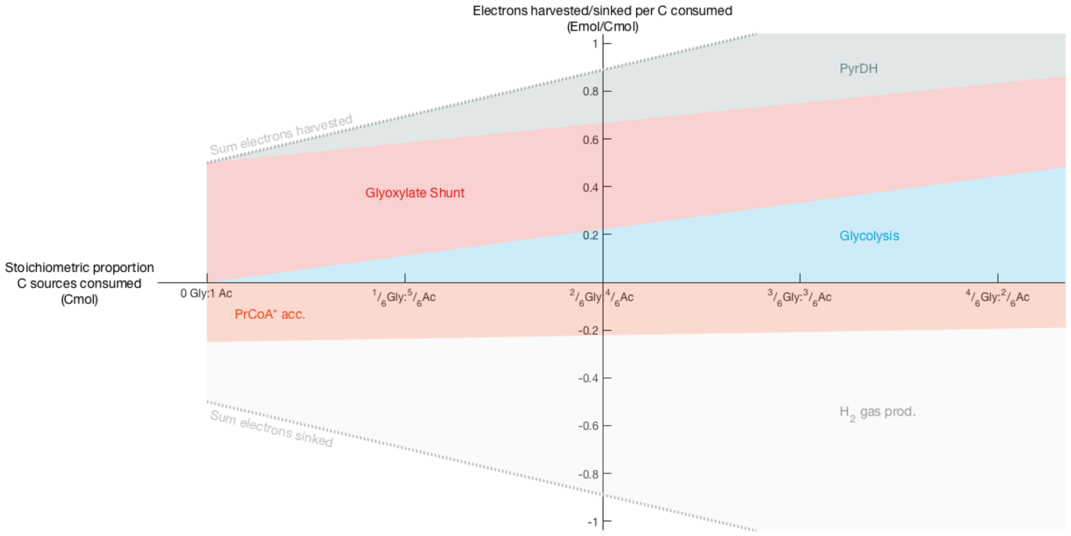


*Figure S6-7 – Electron fluxes through the different sources and sinks depending on the proportion of glycogen to acetate consumed. Values are normalized to C consumed.* ***Optimum scenario*** *– highest Ac-CoA* accumulation;* ***Minimum scenario*** *– lowest carbon conservation.*

##### viii. Alternative scenarios: Full TCA operation?

The full operation of TCA cycle anaerobically in Accumulibacter has been proposed several times, but never convincingly proven. To simulate this, we added the TCA cycle reaction to the stoichiometric matrix, as follows:

TCA: Ac-CoA -> 2$CO_2$ + 8 electrons

The optimum scenario was obtained with the goal of maximum carbon conservation and the minimum scenario with the goal of lowest carbon conservation. The results are shown below.

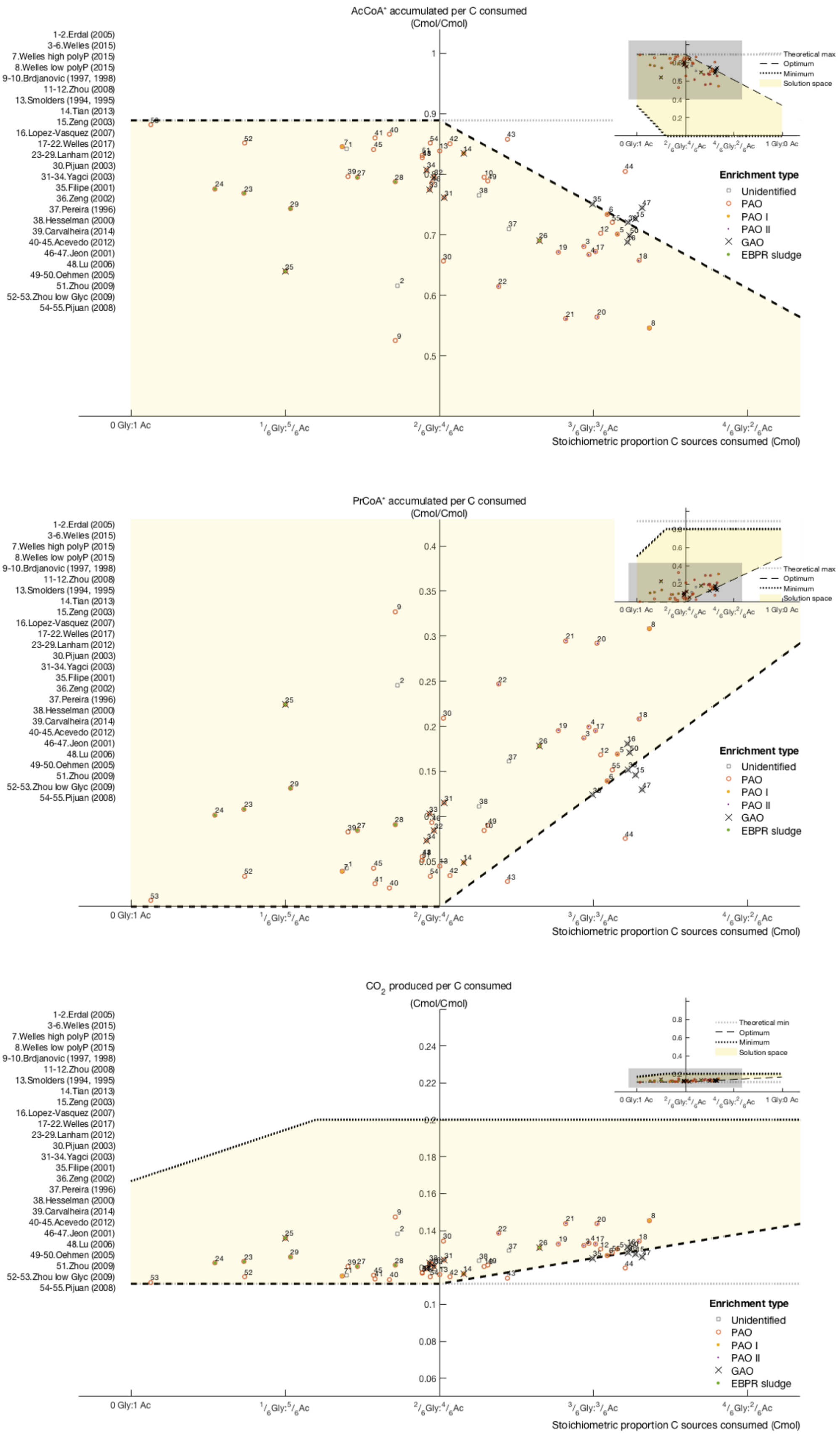


*Figure S6-8 – The full TCA operation allows to explain the amounts of Ac-CoA* and Pr-CoA* found experimentally under the conditions of limitation of glycogen over acetate consumed. Remarkably, this option allows to achieve the maximum theoretical yield of PHB (and PHA) production in "glycogen-limiting" conditions.*

The electron flux distributions can be found below.

Optimum scenario:

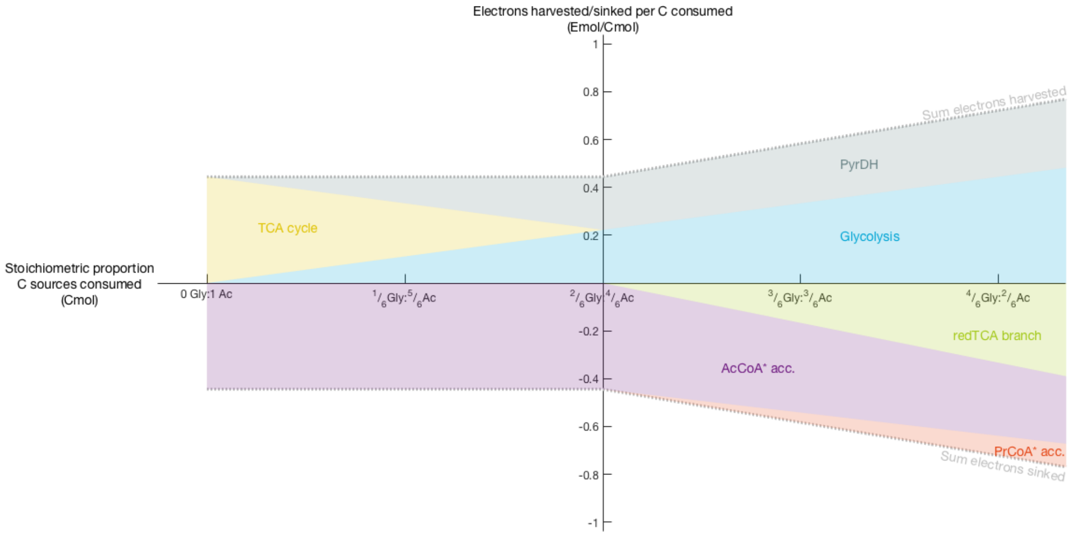


Minimum scenario:

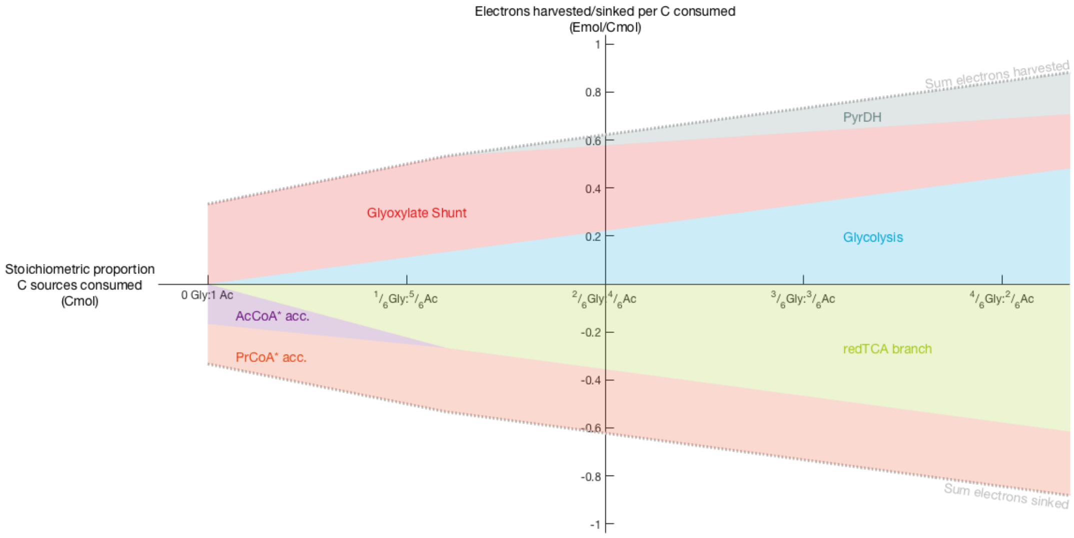


*Figure S6-9 – Electron fluxes through the different sources and sinks depending on the proportion of glycogen to acetate consumed. Values are normalized to C consumed.* ***Optimum scenario*** *– highest carbon conservation;* ***Minimum scenario*** *– lowest carbon conservation.*

#### ix. All scenarios: base case, with $H_2$ production and with full TCA

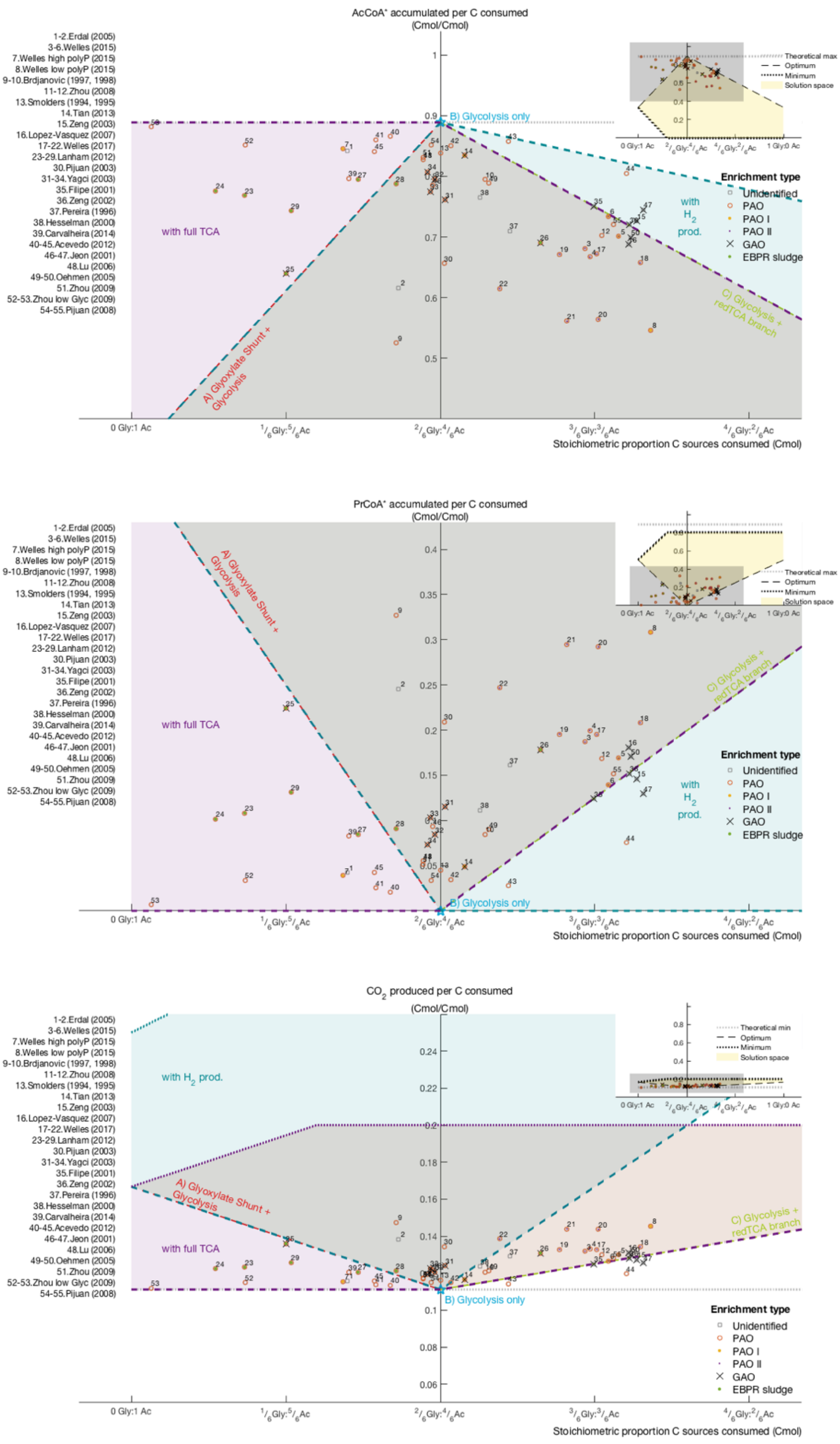


*Figure S6-10 – Overlapped scenarios: Yellow – base case; green – $H_2$ production; purple – full TCA.*
